# Unified Methods for Feature Selection in Large-Scale Genomic Studies with Censored Survival Outcomes

**DOI:** 10.1101/2020.02.14.944314

**Authors:** Lauren Spirko-Burns, Karthik Devarajan

## Abstract

One of the major goals in large-scale genomic studies is to identify genes with a prognostic impact on time-to-event outcomes which provide insight into the disease’s process. With rapid developments in high-throughput genomic technologies in the past two decades, the scientific community is able to monitor the expression levels of tens of thousands of genes and proteins resulting in enormous data sets where the number of genomic features is far greater than the number of subjects. Methods based on univariate Cox regression are often used to select genomic features related to survival outcome; however, the Cox model assumes proportional hazards (PH), which is unlikely to hold for each feature. When applied to genomic features exhibiting some form of non-proportional hazards (NPH), these methods could lead to an under- or over-estimation of the effects. We propose a broad array of marginal screening techniques that aid in feature ranking and selection by accommodating various forms of NPH. First, we develop an approach based on Kullback-Leibler information divergence and the Yang-Prentice model that includes methods for the PH and proportional odds (PO) models as special cases. Next, we propose *R*^2^ indices for the PH and PO models that can be interpreted in terms of explained randomness. Lastly, we propose a generalized pseudo-*R*^2^ measure that includes PH, PO, crossing hazards and crossing odds models as special cases and can be interpreted as the percentage of separability between subjects experiencing the event and not experiencing the event according to feature expression. We evaluate the performance of our measures using extensive simulation studies and publicly available data sets in cancer genomics. We demonstrate that the proposed methods successfully address the issue of NPH in genomic feature selection and outperform existing methods. The proposed information divergence, *R*^2^ and pseudo-*R*^2^ measures were implemented in R (www.R-project.org) and code is available upon request.

## 1 Introduction

There have been significant developments in high-throughput genomic and related technologies in the past two decades. Examples include microarray technology to measure mRNA and microRNA expression, methylation arrays to quantify DNA methylation, SNP arrays to measure allele-specific expression and DNA copy number variation, next-generation sequencing technologies such as RNA-Seq, ChIP-Seq, etc. for the measurement of digital gene expression as well as mass spectroscopy and nuclear magnetic resonance spectroscopy for the measurement of protein and metabolite expression. With the wealth of data available from large-scale genomic studies, researchers can now attempt to understand and estimate the effects of specific genomic features on various diseases and characteristics associated with those diseases. A genomic feature may represent a gene that codes for a protein or a non-gene-centric element such as a microRNA, CpG site or a particular genomic region of interest. One specific area of interest is in studying the relationship between the expression of genomic features and time to death or recurrence of some disease, often referred to as “survival time”. Let *Y* and *C* denote, respectively, the survival and censoring times, and let *δ* = *I*(*Y* ≤ *C*) be the indicator of whether the event has been observed. Because of censoring, it is not possible to observe all true survival times, so we let *T* = min(*Y, C*) be the observed survival time. In this study, we will deal with *p* features measured for each of *n* subjects, where *p* ≫ *n*. We let **Z** denote the *n* × *p* matrix of features and z represent the *p*-vector of features for a subject.

These high-dimensional data sets offer some explicit challenges when applying standard statistical methods. One of the most commonly used models in survival analysis is the Cox proportional hazards (PH) regression model (Cox, 1972) which postulates that the risk (or hazard) of death of an individual, given their feature measurement, is simply proportional to their baseline risk in the absence of the feature. It is a multiplicative hazards model that implies constant hazard ratio (HR). In a high-throughput genomic study, for instance, the PH assumption would imply a constant effect of feature expression on survival over the entire period of follow-up in a study. However, this assumption cannot be verified for each feature and it is also unlikely that PH will actually hold for each feature. Moreover, there is empirical evidence indicating that feature expression may not have a multiplicative effect on the hazard and that non-proportional hazards (NPH) can occur when feature effect increases or decreases over time leading to converging or diverging hazards (CH or DH); in some studies, features with DH are found more often than features with CH (Bhattacharjee et al., 2001; Xu et al., 2005; Dunkler et al., 2010; Rouam et al., 2011). It is, therefore, unreasonable to expect the expression levels of the many thousands of features to exhibit PH. In a review of survival analyses published in cancer journals, it was revealed that only 5% of all studies using the Cox PH model actually checked the PH assumption (Altman et al., 1995). Applying the PH model to data that do not support the underlying assumptions may result in inaccurate and sometimes erroneous conclusions. For instance, it could lead to under- or over-estimation of effects for a considerable number of features. Consequently, some features are falsely declared as important in predicting survival and other relevant features are completely missed. Furthermore, if some features exhibit NPH then their HRs, estimated by ignoring their time-dependence, are not comparable to those of features with PH or of features exhibiting different patterns of NPH. Although NPH typically arises from time-dependent effects of features on survival, it could also result from model mis-specification such as from omitting relevant clinical covariates like age at diagnosis and stage of disease. Feature selection methods involving univariate analyses are particularly prone to this problem. The goal of this study is to discuss alternative models and methods that can be used to link large-scale genomic data to a survival outcome, with the ultimate goal of feature selection under different types of hazards that may be present.

This paper is organized as follows. In §2 we survey a variety of well known survival models, many of which are designed to handle non-proportional hazards (NPH), and provide an overview of marginal screening methods that currently exist in the literature. In §3, we use real-life genomic data to motivate the need for these models. Through this analysis, we identify specific models that fit a large proportion of genomic features and allow for NPH. Then in §4, we propose several feature selection methods that do not rely on the PH assumption and, instead, are developed from models capable of handling NPH. These methods are evaluated using extensive simulations and publicly available large-scale data sets in cancer genomics in §5 and §6. Finally, §7 provides a summary and some concluding remarks.

## 2 A brief survey of survival models and existing methods

### 2.1 The PH model and its generalization

The Cox PH model is a semi-parametric regression model proposed by Cox (1972). The hazard rate, *λ*(*t*|**z**), is defined as the instantaneous risk of an event at time *t* for covariate vector **z**, with Λ(*t*|**z**) representing the cumulative hazard function. The model is given by

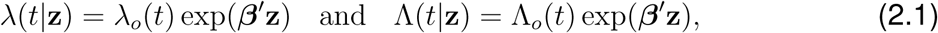

where *t* > 0, *λ*_*o*_(*t*) and Λ_*o*_(*t*) are the baseline hazard and cumulative hazards functions, and ***β*** is a vector of regression coefficients. Estimation for the coefficient ***β*** can be done by maximizing the log partial likelihood 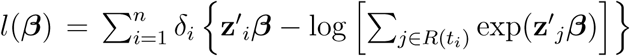, where *t*_*i*_ is the survival time for subject *i, δ*_*i*_ is the censoring indicator, and *R*(*t*_*i*_) is the risk set, the individuals who have yet to experience the event at time *t*_*i*_.

The hazard ratio corresponding to two different feature vectors **z** and **z**^*^, given by 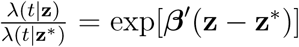, depends only on the features and not on time. This fundamental assumption is unlikely to hold for all *p* features in the genomic setting. A semi-parametric generalization of the Cox PH model which allows crossing hazards is described in Devarajan & Ebrahimi (2011). In this model, the cumulative hazard and survival functions are, respectively,

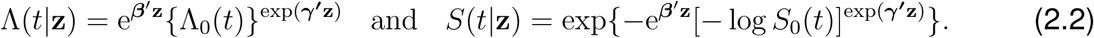

This model has a more flexible, general form, but retains the Cox PH model as a special case when ***γ*** = 0. Since the partial likelihood approach cannot be applied, Devarajan & Ebrahimi (2011) utilize a flexible parametric approach via a cubic *B*-spline approximation for the baseline hazard to estimate the model parameters. Rouam et al. (2011) considered the special case obtained by setting ***β*** = 0 and proposed a pseudo-*R*^2^ measure for genomic feature selection. In this paper, we refer to this special case as the crossing hazards (CH) model.

### 2.2 The proportional odds (PO) model and its generalization

The PO model is an alternative to the PH model, and it does not require the assumption of PH. It allows some forms of NPH and, instead, assumes that the effect of covariates will proportionately increase or decrease the odds of dying or recurrence at time *t*. The PO model is given by

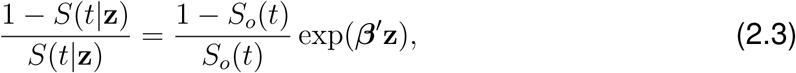

where *S*_*o*_(*t*) and 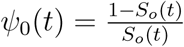 are the baseline survival and odds functions, respectively, at time *t*. The multiplier exp(***β***′**z**) quantifies the amount of proportionate increase or decrease in the odds associated with covariate **z**. A semi-parametric generalization of the PO model can be specified as

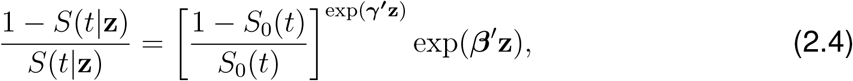

where ***γ*** = 0 results in the PO model. This model allows for both crossing hazards and crossing odds. Later, we consider the special case where ***γ*** = ***β*** and refer to it as the crossing odds (CO) model.

### 2.3 Yang-Prentice (YP) model

Yang & Prentice (2005; 2012) proposed the YP model as a generalization of both the PH and PO models. Its hazard and survival functions are given by

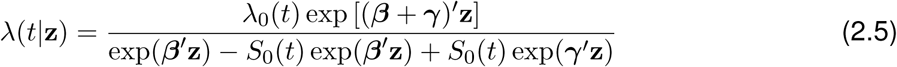

and

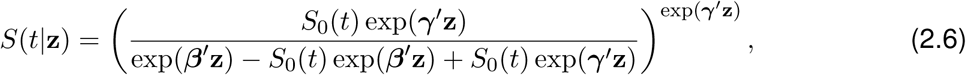

respectively. Note that when ***γ*** = ***β***, it becomes the PH model, and when ***γ*** = 0, it becomes the PO model. Thus, it is a versatile and useful model that encompasses both the PH and PO models, as well as a host of other models, and allows for time-varying hazards. However, a practical limitation of this model is that inferential procedures are available only for a single dichotomized covariate **z**.

### 2.4 Existing methods for feature selection

Few specific methods are currently available in the literature for the purpose of feature selection when the PH assumption is violated. Dunkler et al. (2010) proposed concordance regression, using a generalized concordance probability as a measure of the effect size, to select genes in microarray studies that are related to survival irrespective of the type of hazard. The basic concordance probability is *c* = *P* (*T*_1_ < *T*_0_), where *T*_1_ is a randomly chosen survival time from group 1 and *T*_0_ is a randomly chosen survival time from group 0, and is independent of the PH assumption. This is generalized to continuous data which has the form 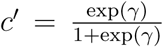, where *γ* are the log odds that the survival time decreases if the gene expression is doubled. Then, *c*′ is modeled through its log odds by 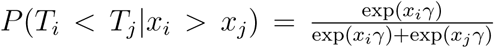, where the likelihood is computed as the summation over all risk pairs (*i, j*) such that *t*_*i*_ < *t*_*j*_ and *c*′ is estimated by maximizing this likelihood. When *t*_*i*_ is censored, it is not clear if *t*_*i*_ < *t*_*j*_ and appropriate weights are used to account for the possible over-representation of some subjects (Schemper et al., 2009). This model can be viewed as conditional logistic regression where the dependent variable is the concordance of the risk pair (*i, j*). Genes are ranked based on the absolute effect size 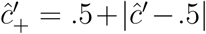. Using simulated and microarray gene expression data Dunkler et al. (2010) showed that when some of the genes showed a time-dependent effect on survival, concordance regression produced the least biased and most stable estimates compared to methods based on Cox regression. Rouam et al. (2010, 2011) developed pseudo-*R*^2^ measures for genomic feature selection based on the PH and CH models which rely on the partial likelihood of the respective model and utilizes the score statistic.

## 3 Motivating examples

We motivate our problem using five data sets from large-scale studies in cancer utilizing a variety of high-throughput genomic technologies. Data sets 1 and 2 consist of DNA methylation and microRNA expression profiles, respectively, measured on glioblastoma samples while data set 3 consists of digital gene expression profiles obtained using RNA sequencing from subjects with head and neck squamous cell carcinoma (The Cancer Genome Atlas (TCGA), http://cancergenome.nih.gov). Data sets 4 and 5 consist of gene expression data obtained using Affymetrix and HG 1.ST microarrays, respectively, from patients with ovarian and oral cancer (Tothill et al, 2008; Saintigny et al, 2011). Raw data was pre-processed using standard methods for each data set as described in the Supplementary Information (SI): Data Sets and Implementation (§8). Relevant characteristics of these data sets are outlined in Tables 1 & 2. In what follows, we describe a comprehensive analysis of these data sets using the PH, PO and YP models to test for the goodness-of-fit (GOF) of each model using the methods outlined in Grambsch & Therneau (1994), Martinussen & Scheike (2006) and Yang & Prentice (2012), respectively. All analyses were performed at the genomic feature level, the purpose of which is to identify features that exhibit some form of NPH and to demonstrate the need for alternatives to the PH model. Wherever possible, clinical covariates such as age at diagnosis and stage of cancer were adjusted for in the analysis (data sets 3-5, Table 1). The currently available method for the YP model is only capable of handling a single dichotomized covariate (Yang & Prentice, 2005; 2012), and, hence, we utilized dichotomized expression of genomic features and did not adjust for age and stage in analyses reported in Table 2. This enabled direct comparison of results from different models. The *q*-value approach was employed to estimate the false discovery rate (FDR) and to evaluate the effect of testing multiple hypotheses (Storey & Tibshirani, 2003). In summary, GOF results based on continuous expression for the PH and PO models (data sets 1-5) as well as dichotomized expression for the PH, PO and YP models (data sets 1, 2 & 4) are presented in Tables 1 and 2, respectively.

**Table 1:**
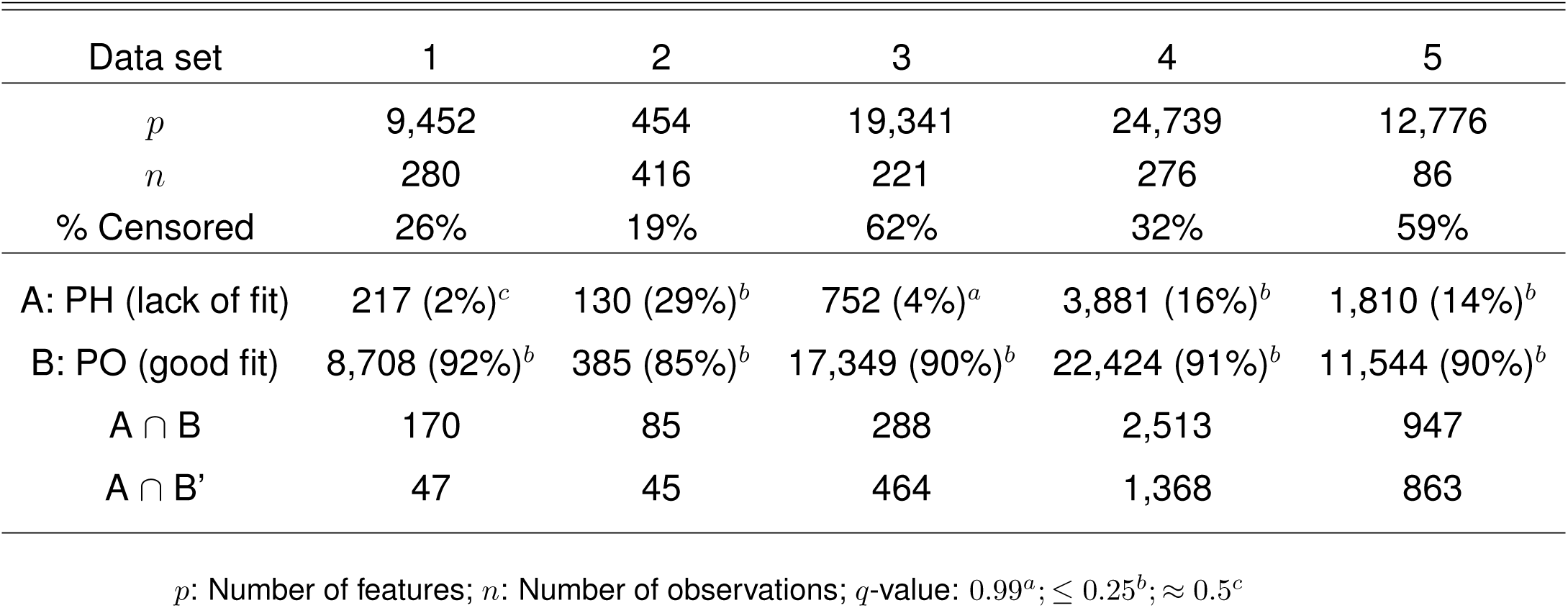
Summary of model fits: continuous data

| Data set | 1 | 2 | 3 | 4 | 5 |
| --- | --- | --- | --- | --- | --- |
| $p$ | 9,452 | 454 | 19,341 | 24,739 | 12,776 |
| $n$ | 280 | 416 | 221 | 276 | 86 |
| % Censored | 26% | 19% | 62% | 32% | 59% |
| A: PH (lack of fit) | 217 (2%)<sup>c</sup> | 130 (29%)<sup>b</sup> | 752 (4%)<sup>a</sup> | 3,881 (16%)<sup>b</sup> | 1,810 (14%)<sup>b</sup> |
| B: PO (good fit) | 8,708 (92%)<sup>b</sup> | 385 (85%)<sup>b</sup> | 17,349 (90%)<sup>b</sup> | 22,424 (91%)<sup>b</sup> | 11,544 (90%)<sup>b</sup> |
| $A \cap B$ | 170 | 85 | 288 | 2,513 | 947 |
| $A \cap B'$ | 47 | 45 | 464 | 1,368 | 863 |

$p$ : Number of features; $n$ : Number of observations; $q$ -value: 0.99<sup>a</sup>; $\leq 0.25$ <sup>b</sup>; $\approx 0.5$ <sup>c</sup>

**Table 2:** Summary of model fits: dichotomized data

| Data set | 1 | 2 | 4 |
| --- | --- | --- | --- |
| $p$ | 9,452 | 454 | 24,739 |
| $n$ | 280 | 416 | 276 |
| % Censored | 26% | 19% | 32% |
| A: PH (lack of fit) | 941 (10%)<sup>b</sup> | 82 (18%)<sup>b</sup> | 3,366 (14%)<sup>b</sup> |
| B: PO (good fit) | 8,880 (94%)<sup>c</sup> | 426 (94%)<sup>b</sup> | 22,628 (91%)<sup>c</sup> |
| C: YP (good fit) | 9,038 (96%)<sup>c</sup> | 441 (97%)<sup>a</sup> | 22,796 (92%)<sup>c</sup> |
| $A \cap B$ | 854 | 72 | 2,325 |
| $A \cap C$ | 896 | 80 | 3,051 |
| $A \cap B' (\cap C)$ | 87 (82) | 10 (9) | 1,041 (926) |

$p$ : Number of features; $n$ : Number of observations; $q$ -value: 0.99<sup>a</sup>; $\leq 0.26$ <sup>b</sup>; 0.33-0.53<sup>c</sup>

Use of continuous expression - typically on a variance stabilized and normalized scale that depends on the data type of the genomic feature - offers a comprehensive approach to our problem and can be used to compute all the *R*^2^-type and information divergence measures developed in this work. It facilitates a straightforward interpretation of the hazard ratio as the fold-change in hazard that corresponds to a unit change in expression on the transformed scale; however, currently it cannot be used for estimation in the YP model and for visualization. On the other hand, dichotomized expression of genomic features - particularly gene expression - is commonly used in real-life applications as evidenced by the literature in high-throughput genomic data analyses (Dunkler et al., 2010; Rouam et al, 2011; Peri et al., 2013). In this approach, the expression of each feature is typically categorized as a “high” or “low” value for each subject based on the median split. Although imperfect, it enables graphical summary of the results in the form of Kaplan-Meier survival curves for subjects with “high” and “low” expression where the hazard ratio is interpreted as the fold-change in hazard between the “low” and “high” expression groups. It can be used for all three models of interest in our problem; however, it is applicable only to information divergence measures.

In Tables 1 and 2, we employ the following abbreviations: Subset A represents genomic features for which the PH model does not fit; B and C refer to subsets of features for which the PO and YP models fit, respectively; and B’ refers to the subset for which the PO model does not fit. As seen from Table 1, the FDR is controlled at an acceptable level (*q*-value ≤ 0.25) in nearly all analyses involving continuous expression. Overall, the PO model is observed to fit a large fraction of genomic features across all data sets (85-92%, subset B). There is clear evidence that the PH model does not fit a significant number of features in these data sets, particularly 2, 4 & 5, where the actual number itself varies across these sets (14-29%, subset A). However, it is important to note that for a majority of the features in these subsets the PO model provides a good fit (52-65%, subset A ∩ B), thereby making this model an attractive alternative to the PH model when continuous expression is considered and potential confounders need to be accounted for. The results for data sets 3, 4 & 5 are further strengthened by an adjustment for confounders such as age at diagnosis and stage of disease. Although the PH model is seen to provide a good fit to most features in data set 1, it is interesting to note that the PO model fits 78% of features (170 out of 217) for which the PH model does not fit. Thus, it would be beneficial to develop methods based on the PO model because it has been shown to handle NPH (Martinussen & Scheike, 2006). In each data set, there exists a reasonable number of features for which neither the PH nor the PO model fits (subset A ∩ B’). This subset contains a median 35% of features for which the PH model does not fit and, in particular, for data set 3 it represents 62% of features (464 out of 752). These observations suggest the need for a more general survival model such as the YP.

As noted earlier, the YP model cannot be used on continuous expression due to current limitation in methods. However, as shown in Table 2, when applied to dichotomized expression the YP model provides a good fit to a majority of the features in each data set considered (92-97%, subset C). This is not surprising given the flexible form of the YP model and its inclusion of the PH and PO models as special cases. More importantly, the PO model is observed to fit a large fraction of genomic features across all data sets (91-94%, subset B). In each data set, the PH model does not fit a large number of features (10-18%, subset A). Once again, the PO model provides a good fit for an overwhelming fraction of features for which PH does not fit - 91%, 88% and 69%, respectively, for data sets 1, 2 and & 4 as shown in Table 2 (subset A ∩ B). In addition, the YP model provides a good fit to an even larger fraction of features for which the PH does not fit - 95%, 98% and 91%, respectively, for these data sets (subset A ∩ C). Furthermore, the YP model serves as a useful alternative for genomic features for which both the PH and PO models do not provide a good fit. In our examples, the YP model provides a good fit for 89-94% of genomic features for which neither PH nor PO fits (subset A ∩ B’ ∩ C). Thus, the YP model shows versatility and the ability to fit a large number of features when the PH and PO models do not. The PO and YP models, therefore, provide flexible alternatives to the PH model when dichotomized expression is considered and it would be useful to develop methods based on these models for handling various forms of NPH. In summary, these results serve as motivation for developing methods based on the PO and YP models because of their ability to fit not only a large number of genomic features in general, but specifically features for which the PH assumption is violated.

## 4 Proposed methods for feature selection and ranking

In §3, we demonstrated the need for alternative methods to the PH model that can handle various types of NPH. We also showed in Tables 1 and 2 that there are many features for which the PH model does not fit but the PO or YP model does; and for some features, we observed that both the PH and PO models do not fit thereby suggesting the need for more complex models such as the YP or CO.

In this section, we construct several marginal screening approaches based on the PO, CO and YP models. First, we adopt an information-theoretic approach and develop a test for genomic feature effect under the YP model using Kullback-Leibler (KL) information divergence. This approach includes tests as well as *R*^2^ measures for its two important special cases - the PO and PH models. Following this, we propose a unified framework to compute pseudo-*R*^2^ measures for a wide range of survival models that allow different types of NPH. It includes the PH, CH, PO and CO models and generalizes prior work (Rouam et al., 2010; 2011). Finally, we propose *R*^2^ measures for the PO model based on the likelihood ratio. The utility of these new marginal screening methods is demonstrated using simulated and real-life genomic data sets and by comparison to existing methods.

### 4.1 Methods based on information divergence

#### 4.1.1 Test for genomic feature effect under the YP model

Within the framework of the YP model, we would like to test the null hypothesis *H*_0_ : Λ(*t*|*z*) = Λ_0_(*t*) against the alternative 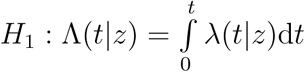, where *λ*(*t*|*z*) is as specified in equation (2.5). These hypotheses can be rewritten as *H*_0_ : *β* = 0, *γ* = 0 vs. *H*_1_ : *β* ≠ 0, *γ* ≠ 0, i.e., we are interested in testing whether a particular genomic feature has an effect on survival time according to the YP model. To this end, we utilize KL information divergence and construct a test statistic. Let *f*_0_(*t*) and *f* (*t*|*z*) denote the densities of *T* under *H*_0_ and *H*_1_, respectively, and let *F*_0_ and *F* be the corresponding distribution functions. Then KL information divergence is defined as

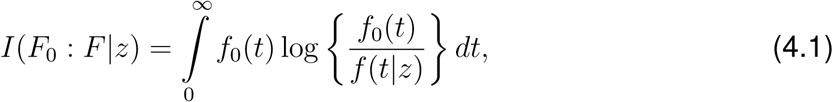

and is the directed divergence that measures the discrepancy between *F*_0_ and *F*. Equation (4.1) quantifies the divergence between the null and alternative hypotheses and can be viewed as a weighted log-likelihood ratio, i.e., 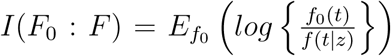. We will use this quantity to develop a test for genomic feature effect.

Under the YP model in equations (2.5) and (2.6), we have *f* (*t*|*z*) = *λ*(*t*|*z*)*S*(*t*|*z*) and *f*_0_(*t*) = *λ*_0_(*t*)*S*_0_(*t*). Hence, feature-specific KL information divergence for the YP model has the form

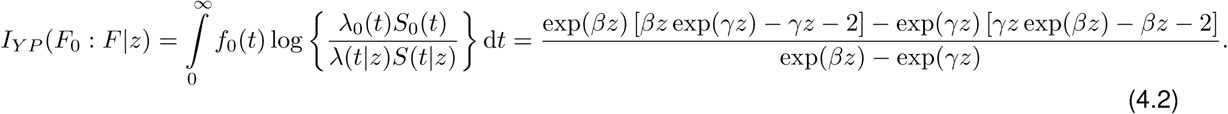

Let *z*_*ij*_ represent the expression for individual *i* and feature *j*, where *i* = 1, …, *n* and *j* = 1, …, *p*. We estimate this measure by replacing *β* and *γ* with 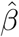 and 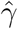, the pseudo maximum likelihood estimates obtained using the approach in Yang & Prentice (2005; 2012), and by summing over the *n* individuals in a study. Thus, we obtain the following estimate of 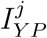 for feature *j*

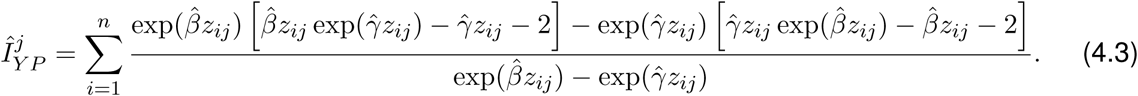

From Theorem 1 (see SI: Methods, §9.1) it can be seen that *Î*_*Y P*_ is a maximum likelihood estimator and is asymptotically normal with mean *I*_*Y P*_. Despite the complexity of the YP model, *Î*_*Y P*_ has the computational advantage of not requiring an estimate of the baseline. In addition, it combines feature effects quantified by the two model parameters *β* and *γ* into a single measure and provides a simpler interpretation of feature effect. However, a practical limitation of the currently available estimation method for this model is that it requires dichotomized feature expression (Yang & Prentice, 2005; 2012). Therefore, we propose *I*_*Y P*_ as a measure for ranking feature effect. In our applications, *I*_*Y P*_ was estimated using 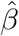 and 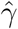 obtained by fitting the YP model to feature expression dichotomized by the median. Details on the derivation of *I*_*Y P*_ are provided in SI: Methods (§9.1).

#### 4.1.2 Tests for genomic feature effect under the PO and PH models

As outlined earlier, the YP model contains the PO and PH models as special cases. In equations (2.5) and (2.6), setting *γ* = *β* results in the PH model and setting *γ* = 0 results in the PO model. In each model there is a single parameter *β* and we are interested evaluating feature effect under the respective model by testing the null hypothesis *H*_0_ : *β* = 0 against the alternative *H*_1_ : *β* ≠ 0.

The test statistic for feature *j* under the PO model is obtained by setting *γ* = 0 in equation (4.2) and using equation (4.3) which simplifies to

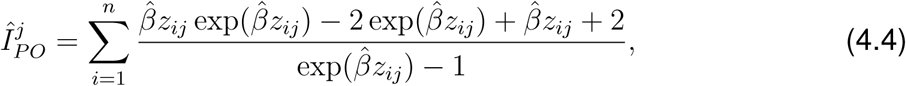

where 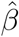 is the modified partial likelihood estimator (Martinussen & Scheike, 2006). From Theorem 2 (see SI: Methods, §9.1, for details on *I*_*PO*_), it can be seen that *Î*_*PO*_ is a maximum likelihood estimator and is asymptotically normal with mean *I*_*PO*_. The variance of *Î*_*P O*_ can be estimated using the delta method and Taylor series expansion as

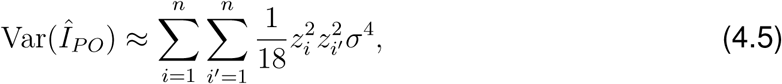

where *σ* is the variance under *H*_0_. Details on the derivation of *V ar*(*Î*_*PO*_) are provided in SI: Methods (§9.1). Using (4.5) we can reject *H*_0_ if

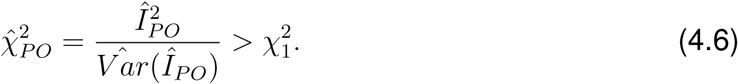

The test statistic for feature *j* under the PH model, *Î*_*PH*_, is obtained by setting *γ* = *β* in equation (4.3) resulting in a test similar to those outlined above for the YP and PO models. This is outlined in Devarajan & Ebrahimi (2009); however, it has not been applied to large-scale genomic data and could potentially over- or under-estimate feature effects when the PH assumption is violated. In that regard, the proposed methods based on YP and PO models benefit from the simplicity of this approach while addressing the issue of NPH.

#### 4.1.3 Feature ranking and selection

Since *Î* quantifies the effect of a genomic feature according to the particular model of interest, it can be directly used to serve as a measure for feature ranking. Both *Î*_*PO*_ and *Î*_*Y P*_ can be calculated for each feature in a data set where a higher *Î* indicates a larger effect on survival based on the particular model chosen. Similarly, the test statistic in equation (4.6) based on the PO model can be used to to compute a *p*-value for each feature and features can be selected by controlling the FDR, at a pre-determined level such as 5%, using the Benjamini-Hochberg approach (Benjamini & Hochberg, 1995). A similar approach can be used for feature selection and ranking using *Î*_*PH*_ under the PH model. It is straightforward to account for potential confounders (such as age and stage of disease) by utilizing the model parameter estimate from the adjusted model in the computation of *Î*_*PO*_ and *Î*_*PH*_ and their respective test statistics. Moreover, standard GOF tests such as those used in §3 can be used to determine which of the three measures to use.

### 4.2 Measures of explained randomness

#### 4.1.2 *R*^2^ measures based on information divergence

We utilize tests for genomic feature effect proposed in §4.1.2 to develop *R*^2^ measures, that quantify the fraction of variation explained, for the PO and PH models. These indices take values on the [0, 1] scale and are easy to interpret. From equation (4.2), feature-specific KL information divergence for the PO model can be obtained by setting *γ* = 0 and expressed as

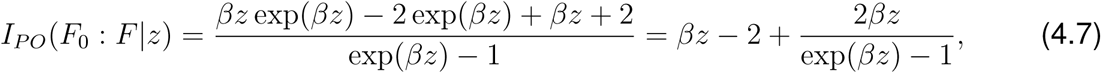

where *z* represents feature expression. Similarly, feature-specific KL information divergence for the PH model can be written as

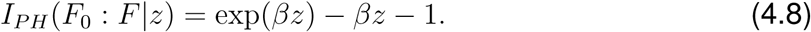

which is obtained in the limit as *β* → *γ* in equation (4.2). In §4.1.1 and §4.1.2, we derived tests for feature effect in the YP and PO models using the respective *Î* by summing over *n* individuals in a data set. This approach accounts for the feature expression of each individual in the study. Here we propose an alternative, but more robust, approach by integrating over the covariate distribution; in this case, the distribution of feature expression. A normalizing transformation stabilizes the variance and can be applied to a variety of large-scale genomic data for this purpose. Examples include the logarithmic transformation for mRNA and miRNA gene expression, the log(*x* + 1) transformation for digital gene expression and the logit transformation for DNA methylation while copy number variation is expressed as log-ratios. In addition, if we standardize the expression of each feature to have zero mean and unit standard deviation, then 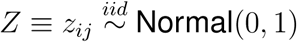 Normal(0, 1), where *i* = 1, …, *n* and *j* = 1, …, *p*. We define *Ĩ* as the expectation of *I*(*F*_0_ : *F*) with respect to the marginal distribution of *Z*,

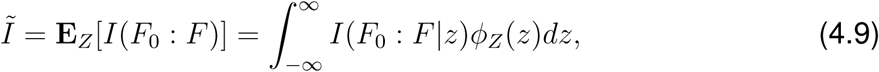

where *ϕ*_*Z*_(*z*) is the standard normal density. *Ĩ* is computed for the PO and PH models using *I*(*F*_0_ : *F* |*z*) in equations (4.7) and (4.8), respectively. An *R*^2^ is then defined as

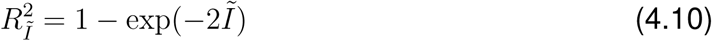

(Joe, 1989; Soofi et al., 1995). For the PO model, *Ĩ*_*PO*_ is calculated using Taylor series expansion as

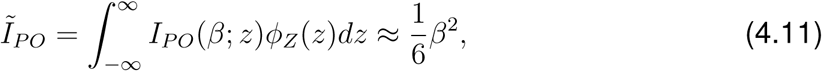

and, hence,

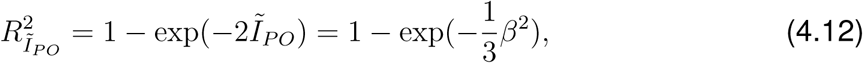

where *β* is replaced by 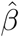, the modified partial likelihood estimator in the PO model obtained using standardized feature expression (Martiussen & Scheike, 2006). For the PH model, we calculate *Ĩ* directly using *I*_*PH*_ as

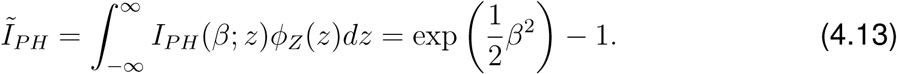

Hence,

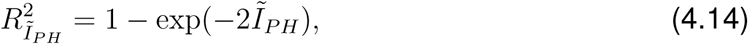

where *β* in *Ĩ*_*PH*_ is replaced by 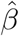, the partial likelihood estimator parameter in the PH model obtained using standardized feature expression (Cox, 1972). Details regarding the derivation of *Ĩ*_*PO*_ and *Ĩ*_*PH*_ are provided in SI: Methods (§9.2). Both 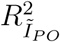 and 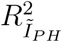 can be easily seen to fall in the [0, 1] range and can be used for feature ranking and selection where genomic features with larger *R*^2^ values can be interpreted as exhibiting larger effects on survival under the respective models. These measures have an information-theoretic foundation and are easy to compute; from equations (4.11) and (4.13), we observe that both measures are simple functions of the respective model parameter *β* which contains the required information if feature expression can be normalized to follow the standard normal model.

#### 4.2.2 *R*^2^ measures based on the likelihood ratio

We propose three different *R*^2^ measures for the PO model based on likelihood ratio (LR), 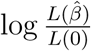, where *L*(0) and 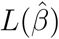 denote the modified partial likelihood for this model under the null and alternative hypotheses, respectively, and the parameter *β* is estimated using the approach outlined in Martinussen & Scheike (2006) (see SI Methods, §9.3, for the modified partial likelihood). These measures parallel corresponding *R*^2^ measures for the PH model that exist in the literature and can be interpreted as the proportion of variation explained by the PO model (Allison, 1995; Nagelkerke, 1991; O’Quigley et al., 2005). The first measure is based on Allison’s index (Allison, 1995) which uses a transformation of the log partial likelihood ratio. It has the form 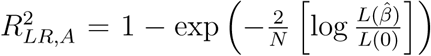 where *N* is the number of subjects. The second measure is a modified version of Allison’s index based on the work in O’Quigley et al. (2005) where *N* in 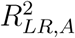 is replaced by *k*, the number of failures. It is less sensitive to censoring which is beneficial in our application due to the observed high fraction of censored observations in large-scale genomic data sets. It is given by

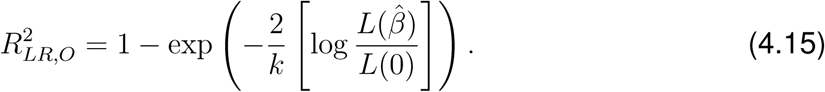

The last measure is based on Nagelkerke’s index (Nagelkerke, 1991) and is another modified version of Allison’s index obtained by dividing the index by its maximum possible value. It has the form 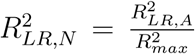 where 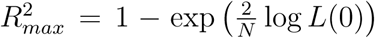. While these measures result in different values and ranges for a specified data set, we note from empirical observation that their rankings are the same. In our simulated study, we found that although their values differed, 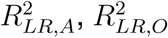, and 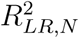 resulted in the same feature rankings. Hence, we choose to use 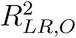 for the remainder of the analysis. For simplicity of notation, we will refer to 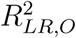 as 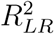 going forward.

### 4.3 A generalized pseudo-*R*^2^ measure

We develop pseudo-*R*^2^ measures for the PO and CO models, denoted by 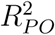 and 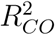, as well as a generalized pseudo-*R*^2^ measure that embeds such measures for the PH, CH, PO and CO models. The proposed approach utilizes the partial likelihood of the respective models and does not require an estimate of the parameter *β* in these models. It generalizes the work of Rouam et al. (2010, 2011) where measures for the PH and CH models, which we denote by 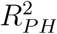 and 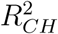, respectively, were proposed. These pseudo-*R*^2^ measures can be interpreted in terms of the difference in the expression of a genomic feature between subjects experiencing and not experiencing the event of interest, and can be used as tools for feature ranking and selection under a variety of scenarios involving NPH. An obvious disadvantage of 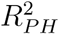 is that it is based on the PH model. In the CH model, the hazard ratio between two individuals with feature expression **z** and **z**^*^ cross over time (Rouam et al., 2011). Thus, while 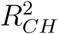 does address the inherent problem with 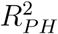, it forces crossing hazards. Therefore, the measure itself is specifically designed to identify crossing hazards. The advantage of the PO model is that it can handle non-proportional hazards while still allowing for proportional hazards, and therefore, is more generally applicable to the variety of hazard structures observed in genomic data. This is evidenced by the results shown in Tables 1 and 2 where PO fits features that exhibit both PH and some forms of NPH. The CO model generalizes the PO model in the same manner as the CH model generalizes the PH model and, thus, has a more versatile form. For our purposes, we will use the special case *γ* = *β* in equation (2.4). Hence, the measures 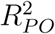 and 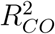 based on these models offer significant advantages and flexibility that is not afforded by currently available measures.

The generalized pseudo-*R*^2^ index can then be expressed as

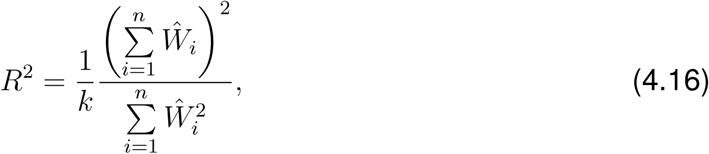

where 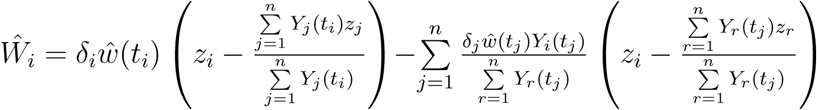, *z*_*i*_ is the expression of a given feature for subject *i, k* is the number of uncensored failure times and 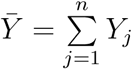 is the number of subjects at risk at time *t*_*i*_ (see SI: Methods, §9.3, for details). This index can be seen as the robust score statistic divided by the number of distinct uncensored failure times, a quantity that falls between 0 and 1 and can be interpreted in terms of the percentage of separability between subjects experiencing and not experiencing the event in relation to the expression of a genomic feature. Using equation (4.16), indices corresponding to the PH, CH, PO and CO models based on model-specific choice of the weight, *w*(*t*), determined by the respective partial likelihood can be obtained (see SI: Methods, §9.3, for a derivation of weights). The estimated weight, *ŵ*(*t*), for each special case is shown in Table 3 and can be interpreted as the derivative of the log hazard ratio for the corresponding model with respect to the parameter *β* evaluated at *β* = 0. In Table 3, Λ_0_(*t*) is estimated by the left-continuous version of Nelson’s estimator and *S*_0_(*t*) is estimated using the Kaplan-Meier (KM) estimator. We note that 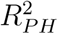 and 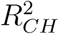 are the measures described in Rouam et al. (2010, 2011) while 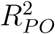 and 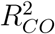 are our newly proposed measures that allow for various types of NPH as well as PH. In order to avoid numerical issues, we applied empirical corrections in the computation of 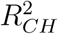 and 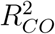 (see SI: Methods, §9.3, for details).

**Table 3:** Pseudo *R*^2^: Special Cases

| Model | Measure | Weight function, $\hat{w}(t)$ |
| --- | --- | --- |
| PH | $R_{PH}^2$ | 1 |
| CH | $R_{CH}^2$ | $1 + \log\{\hat{\Lambda}_0(t)\}$ |
| PO | $R_{PO}^2$ | $\hat{S}_0(t)$ |
| CO | $R_{CO}^2$ | $1 + \hat{S}_0(t) - \hat{S}_0(t) \log\left(\frac{\hat{S}_0(t)}{1 - \hat{S}_0(t)}\right)$ |

## 5 Application to simulated data

In this section, we evaluate our newly proposed ranking methods, *I*_*PO*_, *I*_*Y P*_, 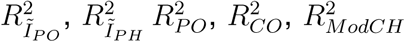 and 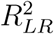 using simulated data sets under a variety of scenarios and compare their performance to existing methods, 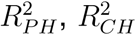 and 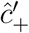 (Rouam et al., 2010, 2011; Dunkler et al., 2010). The goal of this study is to assess the performance of these methods in selecting features that are truly associated with survival in the high-dimensional setting.

### 5.1 Simulation schemes

We considered two different simulation schemes to generate artificial survival and genomic data sets based on the approach outlined in Dunkler *et al.* (2010). In order to account for various types of hazards, survival times *Y*_*i*_, *i* = 1, …, *n*, were generated from each of 5 different models specified as follows: standard log-normal LN (*µ* = 0, *σ* = 1); log-logistic LL1 (*α*_1_ = 2, *λ*_1_ = 2, *λ*_2_ = 4) and LL2 (*α*_1_ = 3, *α*_2_ = 4, *λ*_1_ = 1, *λ*_2_ = 2); and Weibull W1 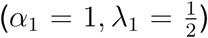 and W2 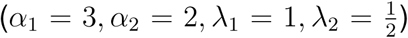, where LL1 and W1 refer to the respective models where the shape parameters are the same but the scale parameters differ, and LL2 and W2 refer to the respective models where both the shape and scale parameters differ. In the LN model, *µ* and *σ* are the location and scale parameters, respectively. We use a more informed approach that is broader in scope compared to that of Dunkler *et al.* (2010), who only considered W1 in their simulations. Here, LN, LL2 and W2 cases are of particular interest because of their ability to simulate crossing hazards. To simulate censoring, we drew random samples with uniform follow-up times *C* from *U* (0, *τ*) and defined the observed survival time as *T* = min(*Y, C*) with censoring indicator *δ* = *I*(*T* = *Y*). We chose *τ* to get censoring proportions of 0, 33, and 67%.

For each model, we simulated censored survival times and genomic data for *N* = 200 subjects and *p* = 5000 mock features whose expression is linked to survival time based on the logarithm of the hazard ratio (HR), *β*_*g*_(*t*) = *β*_0_ log(*HR*). Genomic data was generated from the standard normal model which covers a variety of features seen in large-scale genomic studies. Following Klein and Moeschberger (2003), log(*HR*) was calculated based on the respective model of interest. For LN, we used *β*_*g*_(*t*) = *β*_0_(*t*^2^ − 1) to simulate crossing hazards similar to what was done in Dunkler *et al.* (2010). Then, *β*_0_ was chosen so that only the first 400 features were assumed to have an effect on survival time, with 200 having a large effect and 200 having a small effect. In Scheme 1, we adopt a univariate approach where the expression of each feature is separately linked to survival, and in Scheme 2 we adopt a multivariate approach that incorporates correlations between features. More details on these steps can be found in Dunkler *et al.* (2010).

For each simulation scheme and censoring combination, 200 data sets were generated and assessed. The ranking methods developed §4 were applied to each data set and genomic features were ranked based on each method. The results are summarized using mean AUC, specificity, sensitivity and the Youden index (Youden, 1950) across the 200 simulations in each case and used to compare the methods. The Youden index is calculated as *J* = *sensitivity* + *specificity* − 1 where higher values are desirable.

### 5.2 Simulation Results

Overall, under different models (LN, LL1, LL2, W1, W2) and censoring proportions (0, 33, 67%), the proposed methods outperformed and, in some cases, performed as well as existing methods. In most cases, we noticed some form of improvement. Detailed simulation results under various scenarios outlined above are provided in SI: Simulation Results and SI: Tables 5-9 (§10). Our PO model-based methods performed strongly overall and *I*_*Y P*_ performed particularly well for lower censoring. 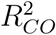 performed similarly to 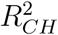 in many cases, but it is important to note that our modified version, 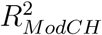, performed better than 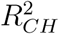 in most cases and similarly to it in other cases. Overall, depending on the simulation scheme and type of non-proportional hazards present, we can identify the benefits of utilizing each of our measures. Most importantly, the proposed feature selection methods are more flexible and generalize existing methods.

**Table 4:**
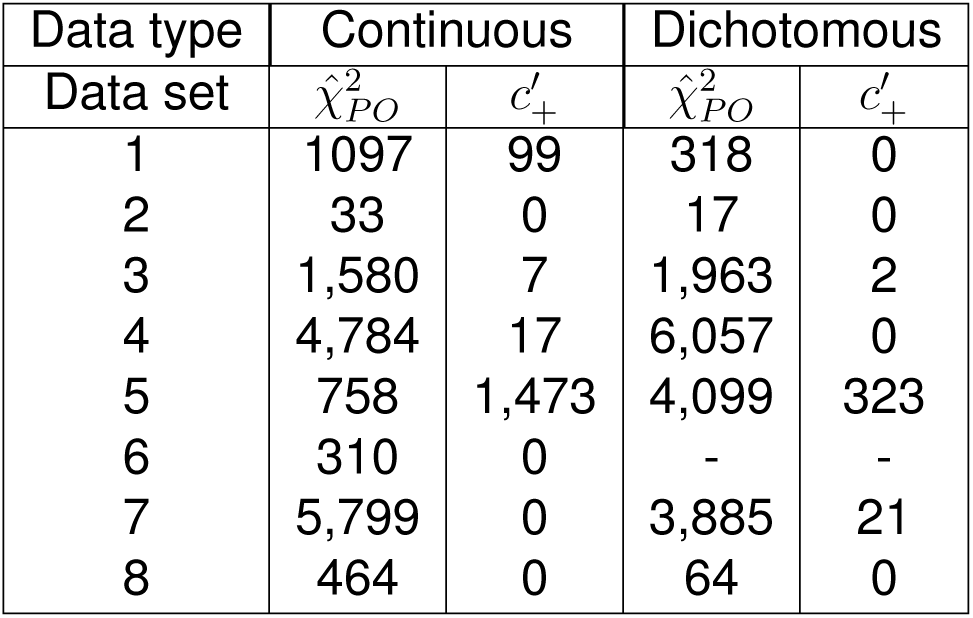
Number of features selected by 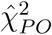 and 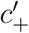

**Table 5:** Simulation scheme 1: Comparison of methods (AUC)

| Censoring | Measure | LN | LL1 | LL2 | W1 | W2 |
| --- | --- | --- | --- | --- | --- | --- |
| 0% | $I_{PO}$ | 0.81 | 1.00 | 0.74 | 1.00 | 1.00 |
| | $I_{YP}$ | 1.00 | 0.96 | 0.81 | 1.00 | 1.00 |
| | $\hat{c}'_+$ | 0.81 | 1.00 | 0.73 | 1.00 | 1.00 |
| | $R^2_{PO}$ | 0.81 | 1.00 | 0.73 | 1.00 | 1.00 |
| | $R^2_{LR}$ | 0.81 | 1.00 | 0.73 | 1.00 | 1.00 |
| | $R^2_{I_{PO}}$ | 0.81 | 1.00 | 0.75 | 1.00 | 1.00 |
| | $R^2_{CO}$ | 0.99 | 0.86 | 0.84 | 0.93 | 1.00 |
| | $R^2_{CH}$ | 1.00 | 0.68 | 0.87 | 0.65 | 0.98 |
| | $R^2_{ModCH}$ | 1.00 | 0.73 | 0.89 | 0.69 | 0.98 |
| | $R^2_{I_{PH}}$ | 0.92 | 0.99 | 0.54 | 1.00 | 1.00 |
| | $R^2_{PH}$ | 0.87 | 0.99 | 0.50 | 1.00 | 1.00 |
| 33% | $I_{PO}$ | 0.89 | 0.99 | 0.77 | 0.99 | 1.00 |
| | $I_{YP}$ | 0.96 | 0.89 | 0.75 | 0.98 | 1.00 |
| | $\hat{c}'_+$ | 0.82 | 0.99 | 0.71 | 0.99 | 1.00 |
| | $R^2_{PO}$ | 0.89 | 0.99 | 0.77 | 0.99 | 0.99 |
| | $R^2_{LR}$ | 0.88 | 0.99 | 0.77 | 1.00 | 1.00 |
| | $R^2_{I_{PO}}$ | 0.88 | 0.99 | 0.78 | 1.00 | 1.00 |
| | $R^2_{CO}$ | 0.82 | 0.69 | 0.80 | 0.82 | 0.87 |
| | $R^2_{CH}$ | 0.98 | 0.82 | 0.86 | 0.70 | 0.76 |
| | $R^2_{ModCH}$ | 0.97 | 0.84 | 0.87 | 0.74 | 0.82 |
| | $R^2_{I_{PH}}$ | 0.77 | 0.99 | 0.64 | 1.00 | 1.00 |
| | $R^2_{PH}$ | 0.77 | 0.99 | 0.61 | 1.00 | 1.00 |
| 67% | $I_{PO}$ | 0.93 | 0.98 | 0.81 | 0.99 | 0.95 |
| | $I_{YP}$ | 0.83 | 0.70 | 0.58 | 0.81 | 0.93 |
| | $\hat{c}'_+$ | 0.88 | 0.96 | 0.71 | 0.98 | 0.98 |
| | $R^2_{PO}$ | 0.93 | 0.97 | 0.81 | 0.99 | 0.95 |
| | $R^2_{LR}$ | 0.93 | 0.98 | 0.81 | 0.99 | 0.95 |
| | $R^2_{I_{PO}}$ | 0.93 | 0.98 | 0.81 | 0.99 | 0.95 |
| | $R^2_{CO}$ | 0.68 | 0.63 | 0.74 | 0.51 | 0.89 |
| | $R^2_{CH}$ | 0.88 | 0.90 | 0.83 | 0.87 | 0.57 |
| | $R^2_{ModCH}$ | 0.87 | 0.90 | 0.84 | 0.87 | 0.71 |
| | $R^2_{I_{PH}}$ | 0.92 | 0.98 | 0.78 | 0.99 | 0.97 |
| | $R^2_{PH}$ | 0.92 | 0.97 | 0.78 | 0.99 | 0.97 |

## 6 Application to genomic data

In this section, we compare the performance of the proposed methods using several data sets representing a broad spectrum of high-throughput genomic technologies. In addition to data sets 1-5 described in §3, we utilized the following data sets. Data set 6 consists of microarray gene expression profiles measured on the same set of glioblastoma samples used in data sets 1 and 2, while data sets 7 and 8 consist of DNA methylation and copy number variation profiles, respectively, from subjects with head and neck squamous cell carcinoma. Since we do not have prior knowledge on the number of genomic features significantly associated with survival in real data, this approach will differ from that of the simulations. Here, we will rank features based on each method and compare them using a preselected number of top features across methods. Whenever possible, parameter estimates required for computing certain *I* and *R*^2^ measures were obtained from respective models adjusted for potential confounders such as age and stage of disease. These measures include *I*_*P O*_, 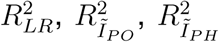 and the absolute effect size estimate, 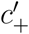, from concordance regression. In addition to continuous feature expression, dichotomized expression was utilized for *I*_*PO*_ and 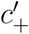 and no adjustment for confounders was done for analyses involving dichotomized data in order to enable direct comparison of results to *I*_*Y P*_ which accommodates only a single dichotomized covariate due to current limitations in the YP model implementation (Yang & Prentice, 2005; 2012). Overall, the approach outlined above for computing the proposed *I* and *R*^2^ measures parallels the analyses presented in §3 comparing different models.

We begin by focusing attention on the application of our proposed *I* measures - *I*_*PO*_ and *I*_*Y P*_ - and 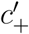, selecting the top 500 features based on each measure. We observe that there are few features commonly selected by all three measures, as evidenced in the Venn diagrams presented in SI: Figures 1 (data sets 1, 2 and 6) and 2 (data sets 3, 7, 8, 4, and 5). This is not surprising given that these measures are based on different model assumptions. However, it is interesting to note that in the glioblastoma data sets (1, 2 and 6) and in the HNSCC data sets 7 and 8, a large fraction of the selected features (62%, 88%, 75%, 62% and 53% respectively) are common to *I*_*PO*_ and 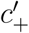. Overall, relatively fewer features appear to be commonly selected by *I*_*Y P*_ and 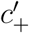. For dichotomized data, it is evident that feature sets corresponding to different measures are much less concordant compared to continuous expression, as shown in SI: Figure 3 (data sets 1, 2 and 4).

**Figure 1:**
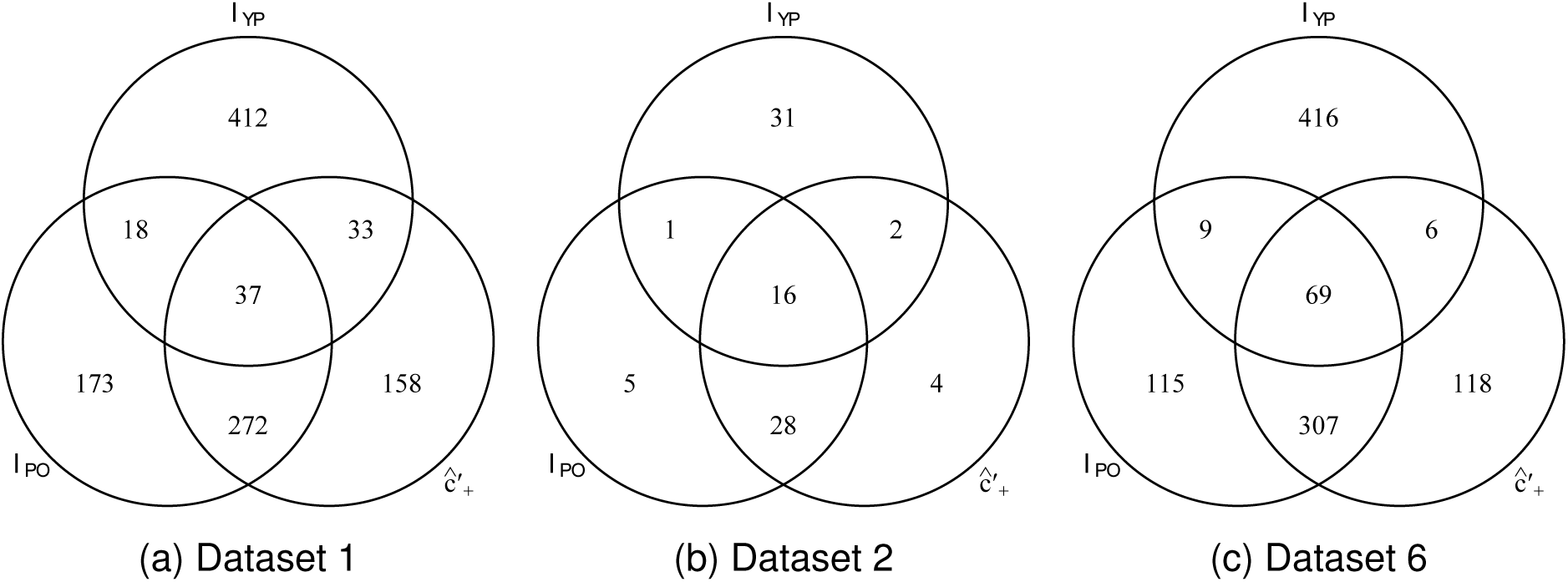
Top selected genes, *I* measures and 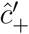, Glioblastoma data sets

**Figure 2:**
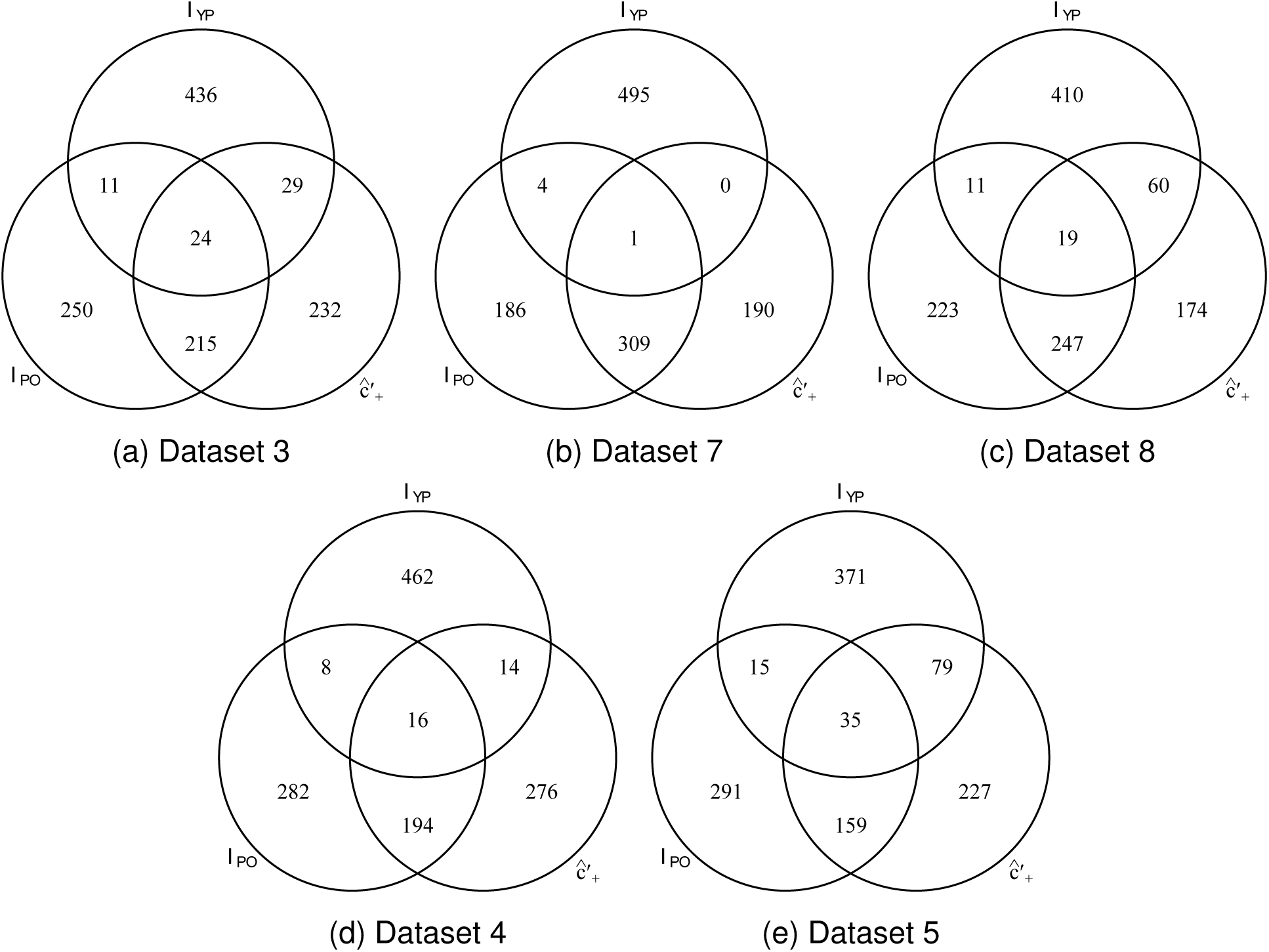
Top selected genes, *I* measures and 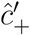, HNSCC (3, 7, and 8), ovarian (4) and oral (5) data sets

**Figure 3:**
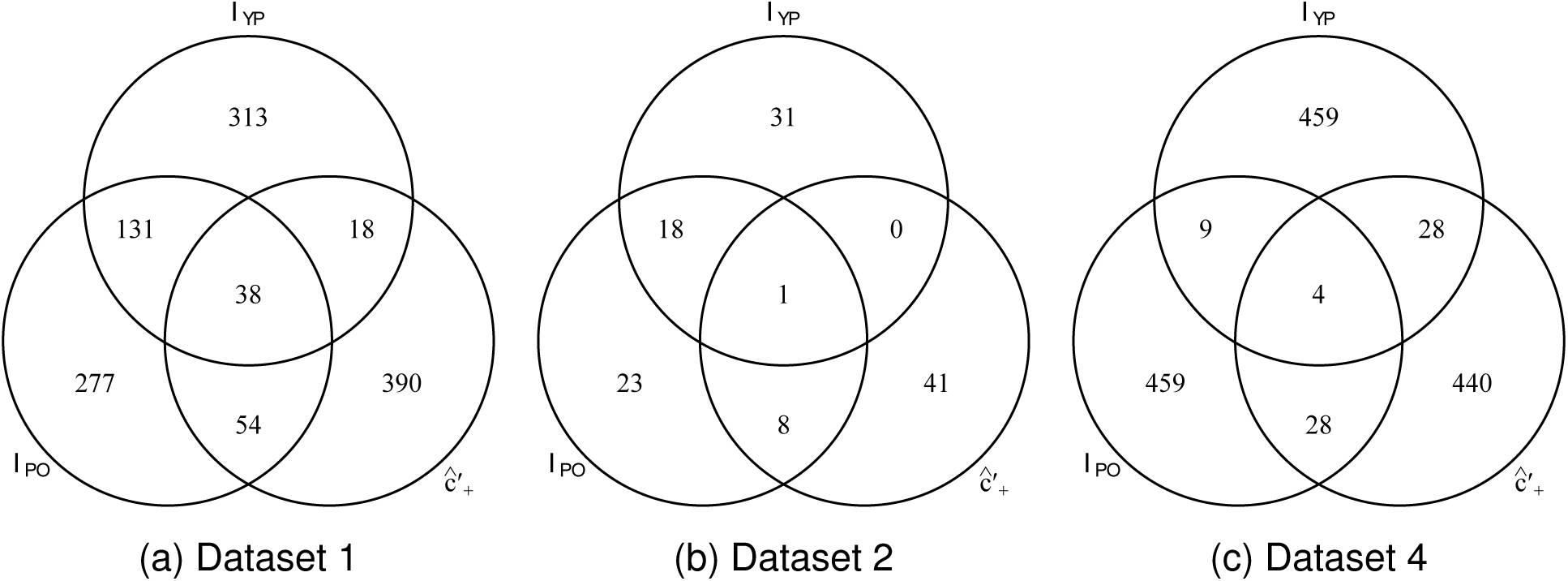
Top selected genes, Dichotomized feature expression, Data sets 1, 2 and 4

Using the test statistic for *Î*_*PO*_ developed in §4 and denoted by 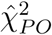, we computed a *p*-value for each feature and selected features by controlling the FDR at 5% using the Benjamini-Hochberg approach (Benjamini & Hochberg, 1995). It should be noted that this approach can also be used to rank features and to compare different measures. Table 4 summarizes the results of this analysis for continuous and dichotomized feature expression for various data sets. In almost all cases, 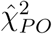 selects a much larger number of features compared to 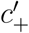 which does not select many features at all in the first place. This is likely due to its more general form which accounts for PH as well some form of NPH. Although adjustment for confounding effects of age and stage or for multiple testing typically reduce the number of features significantly associated with survival, we observed that 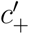 identified fewer features even without such adjustments and fared poorly overall compared to 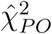 not only in selecting features with some type of NPH but also in selecting features with PH, based on statistical significance. This limitation is recognized in Dunkler et al. (2010) and it renders 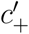 as a tool for feature ranking only rather than feature selection, unlike 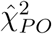 which can both be used for ranking as well as selecting features based on a pre-specified *p*-value or FDR threshold. These observations provide an argument in favor of the use of *I*_*PO*_, 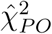 for feature selection and ranking. In SI: Application to Genomic Data (§11) and SI: Figures 8-12, we outline an approach to evaluate and visualize the combined effect of features selected on survival. We thus recommend the use of *I*_*PO*_, 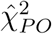 or *I*_*Y P*_ because of their inherent versatility. While *I*_*Y P*_ is able to handle various types of hazards and retains both *I*_*PO*_ and *I*_*PH*_ as special cases, its performance could be significantly improved by developing methods to estimate the YP model parameters for continuous data which are currently unavailable. On the other hand, *I*_*PO*_ performed better than or at least similarly to 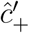 in every simulation scenario considered. All three measures are easy to calculate and the associated statistical test can be used for simple feature selection at a pre-defined significance or FDR threshold, as shown in the examples in this section.

**Figure 4:**
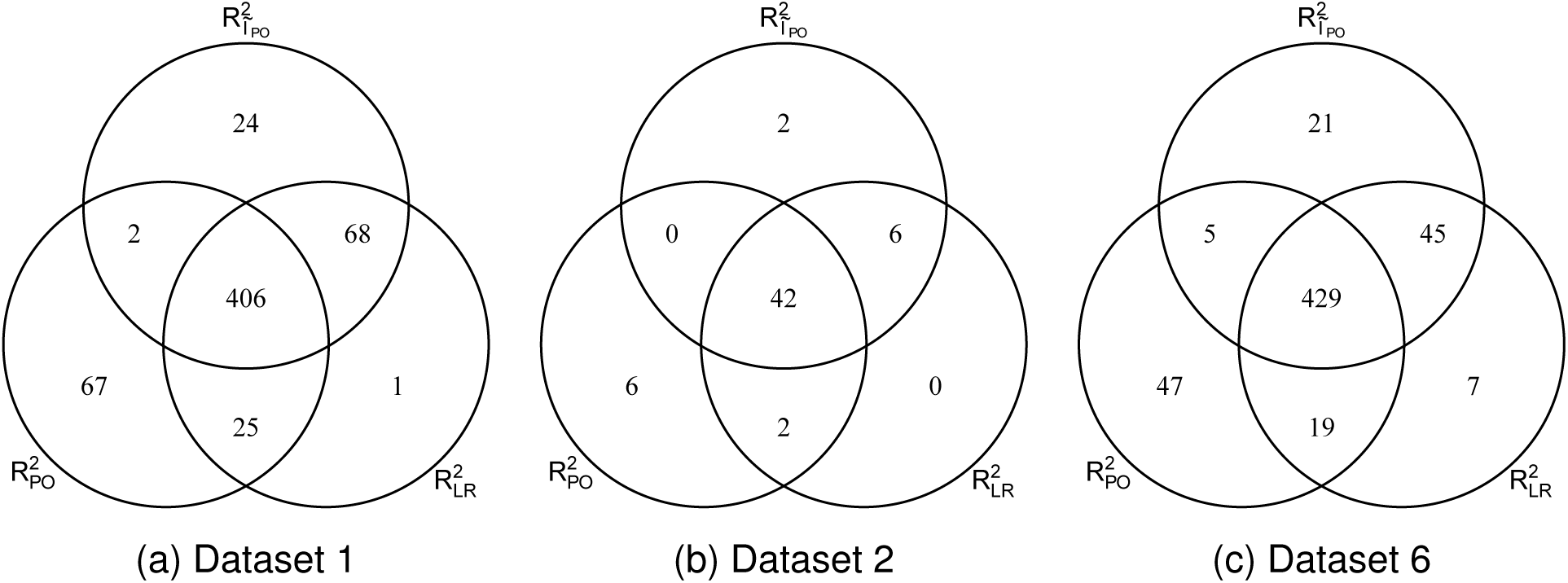
Top selected genes, PO-based *R*^2^ measures, Glioblastoma data sets

**Figure 5:**
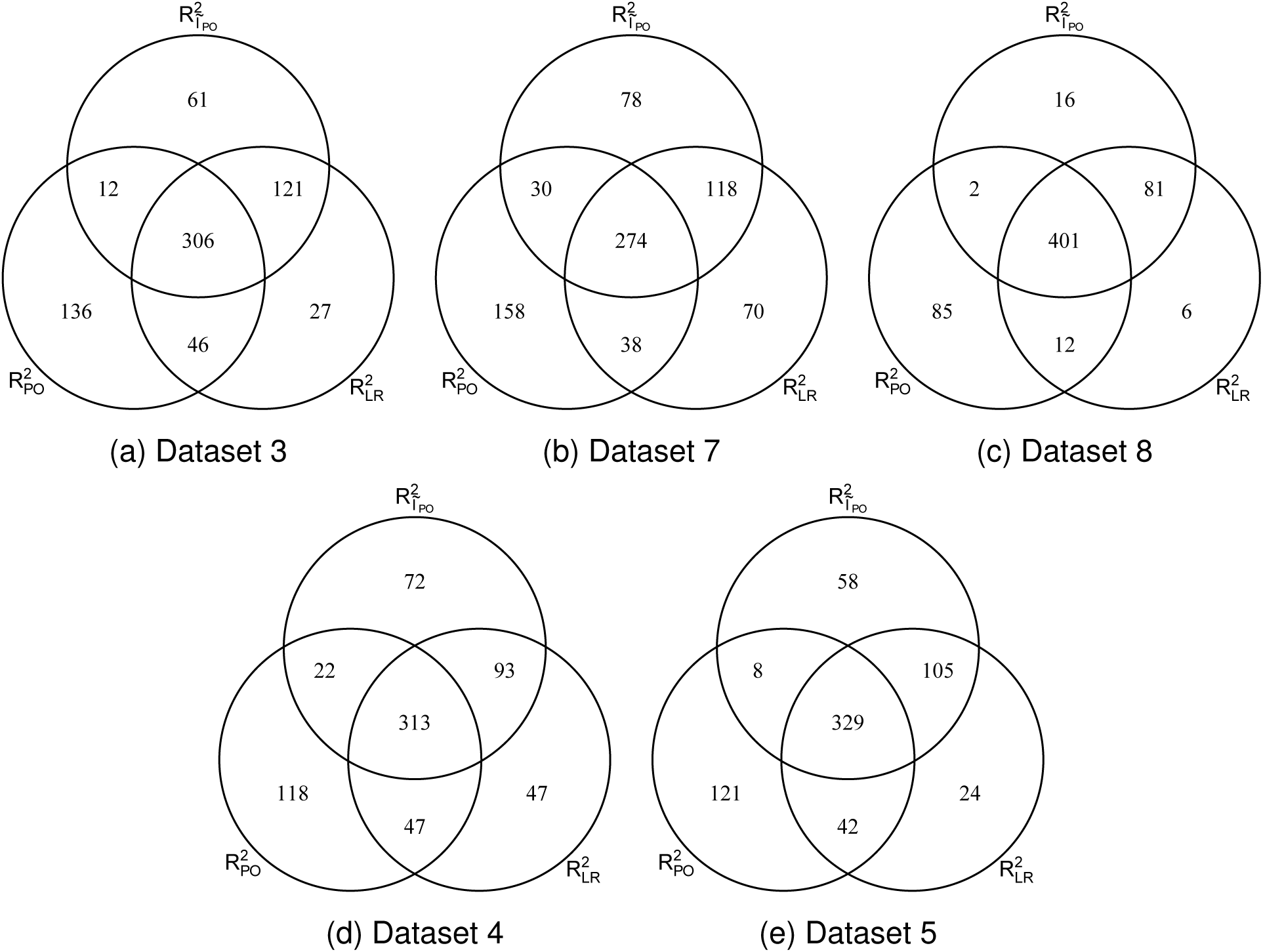
Top selected genes, PO-based *R*^2^ measures, HNSCC (3, 7, and 8), ovarian (4) and oral (5) data sets

**Figure 6:**
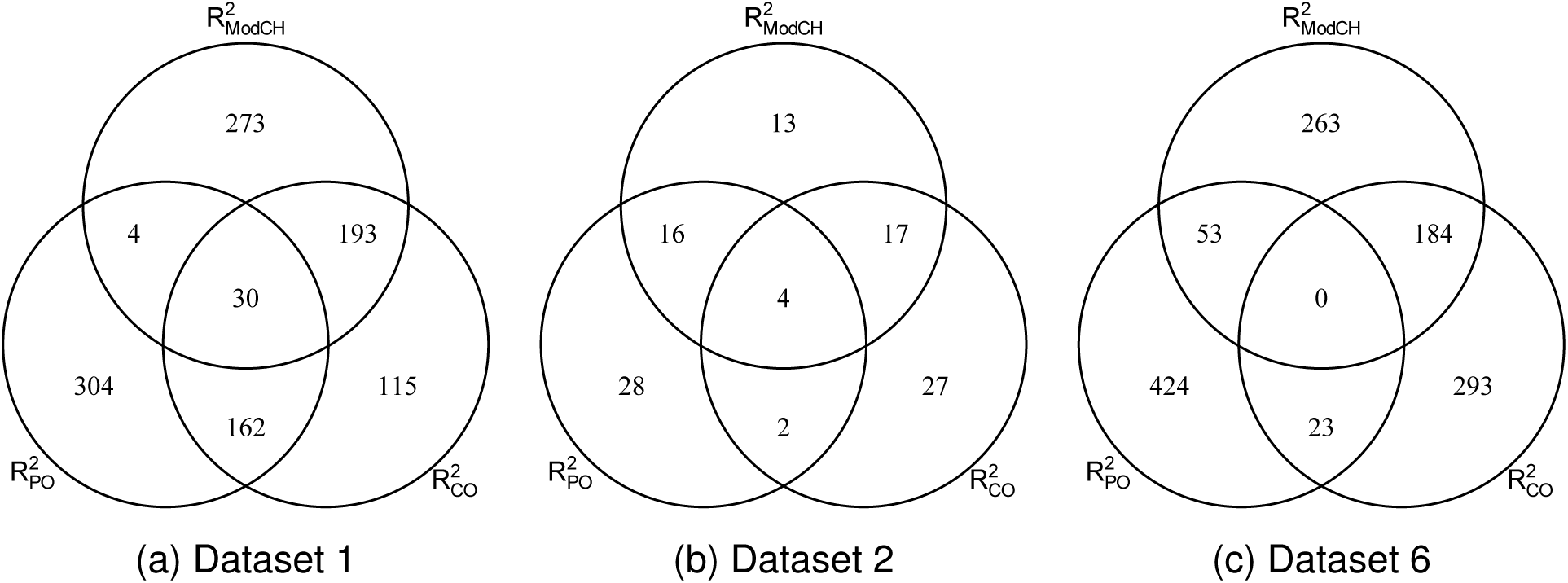
Top selected genes, Other *R*^2^ measures, Glioblastoma data sets

**Figure 7:**
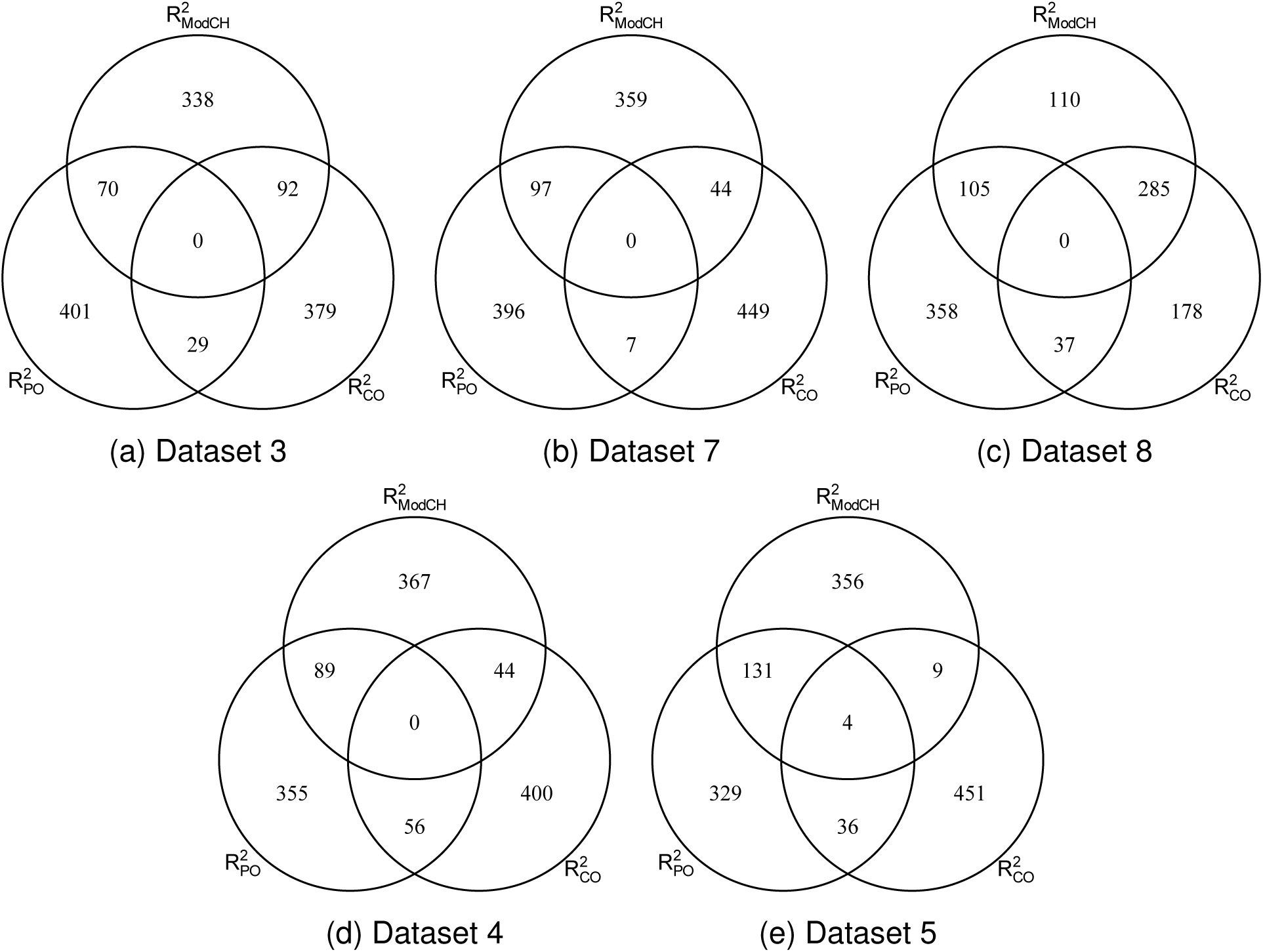
Top selected genes, Other *R*^2^ measures, HNSCC (3, 7, and 8), ovarian (4) and oral (5) data sets

**Figure 8:**
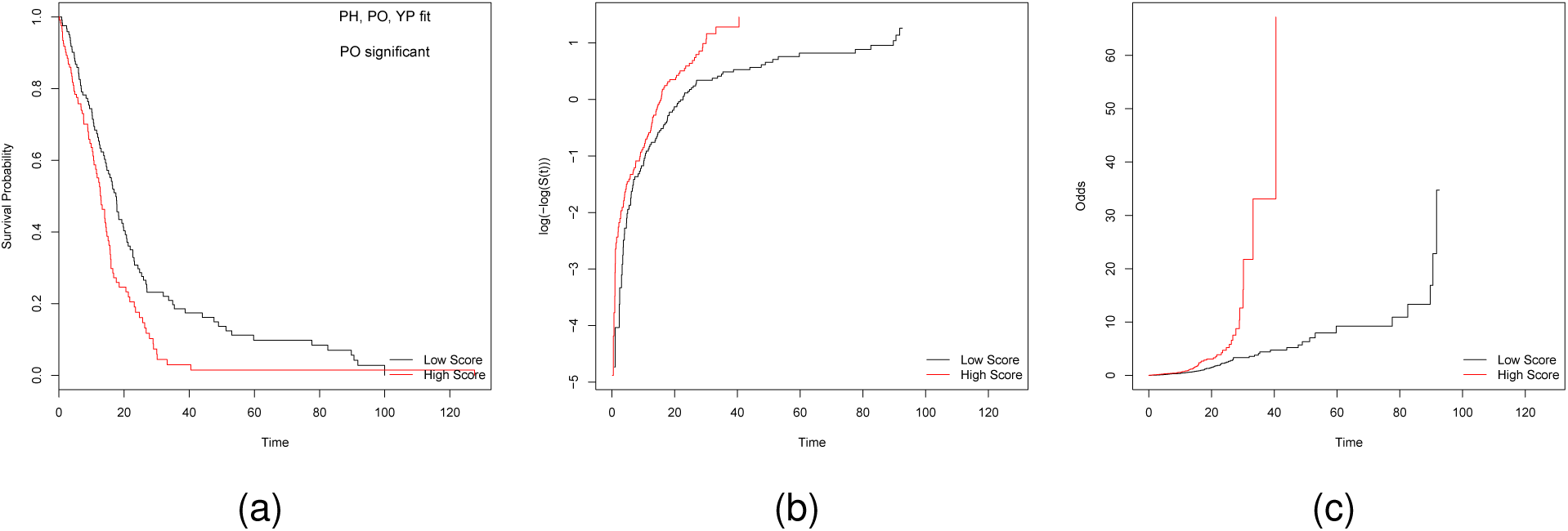
(a) Kaplan-Meier survival curves, (b) Cumulative hazard curves (log-scale), (c) Odds curves. Weighted expression of features selected by 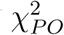 for data set 1

**Figure 9:**
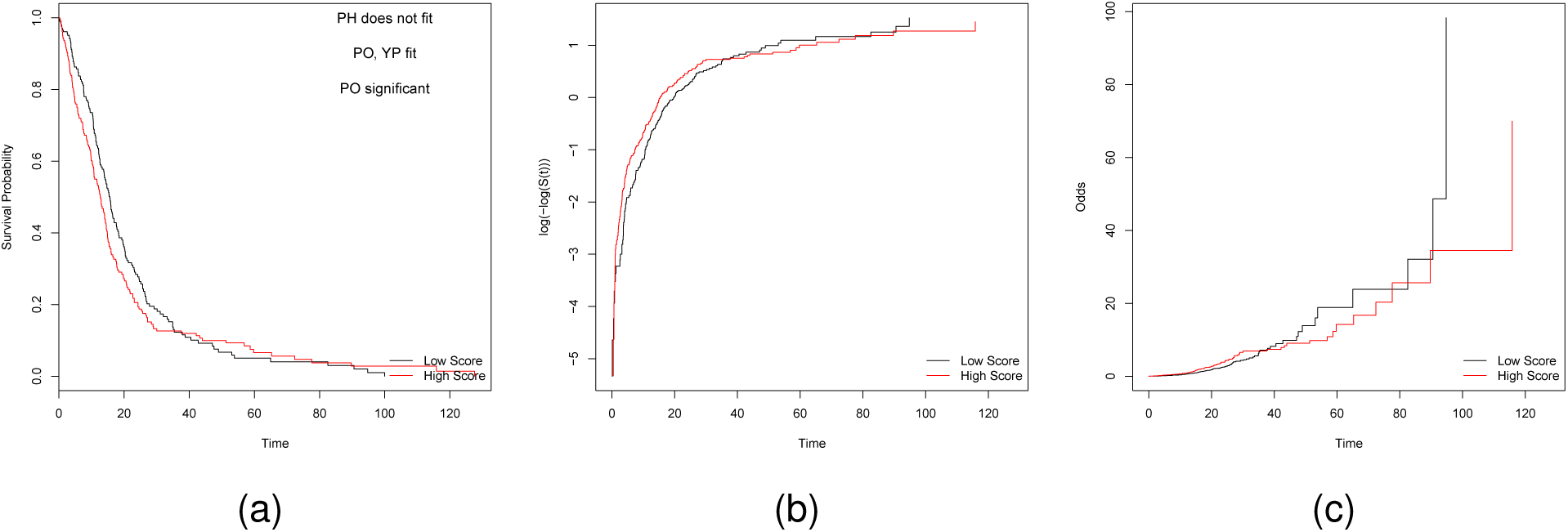
(a) Kaplan-Meier survival curves, (b) Cumulative hazard curves (log-scale), (c) Odds curves. Weighted expression of features selected by 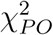 for data set 2

**Figure 10:**
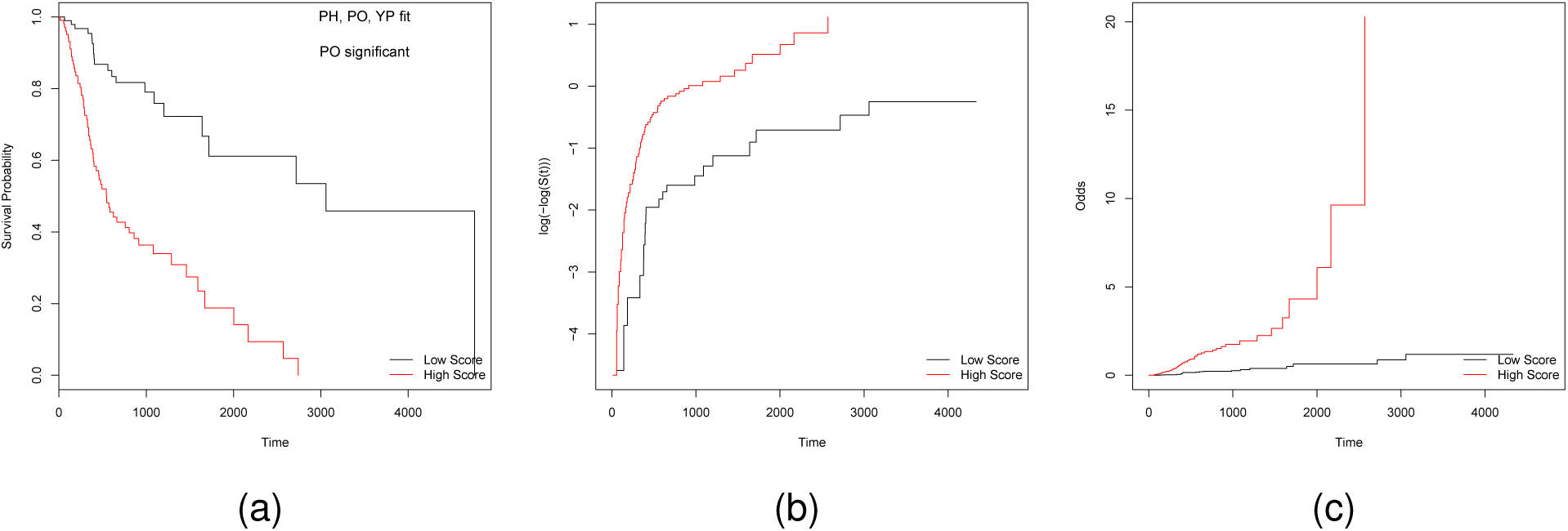
(a) Kaplan-Meier survival curves, (b) Cumulative hazard curves (log-scale), Odds curves. Weighted expression of features selected by 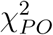 for data set 3

**Figure 11:**
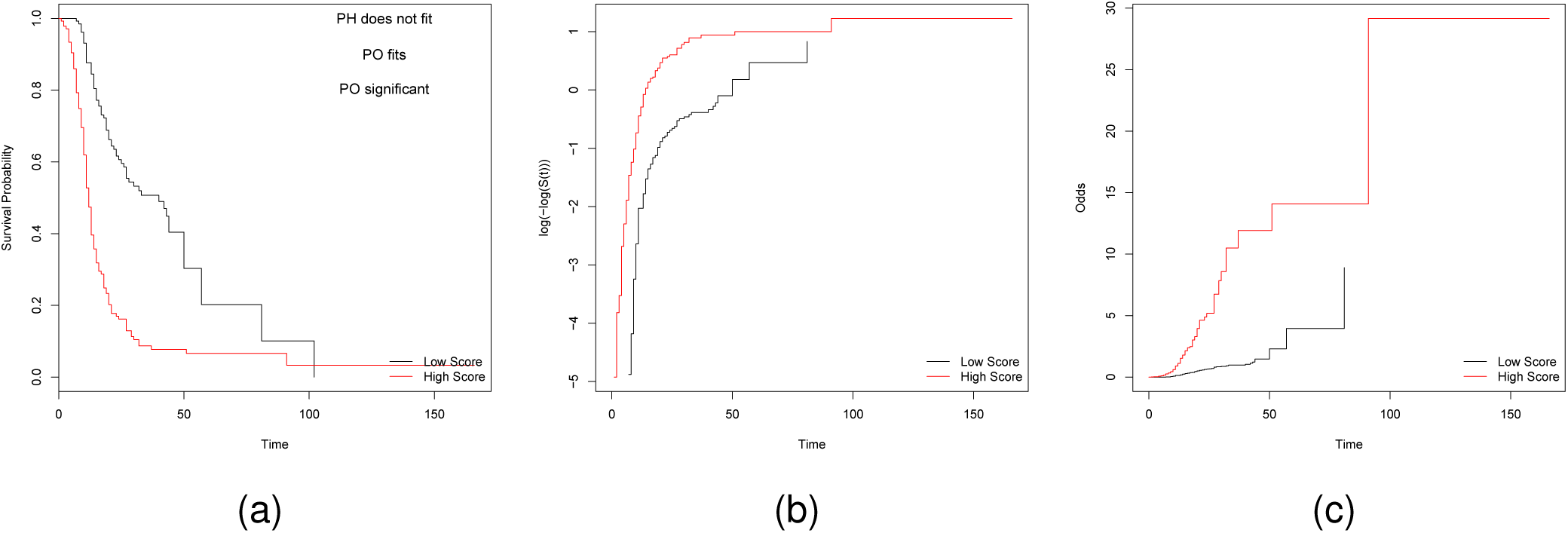
(a) Kaplan-Meier survival curves, (b) Cumulative hazard curves (log-scale), Odds curves. Weighted expression of features selected by 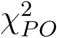 for data set 4

**Figure 12:**
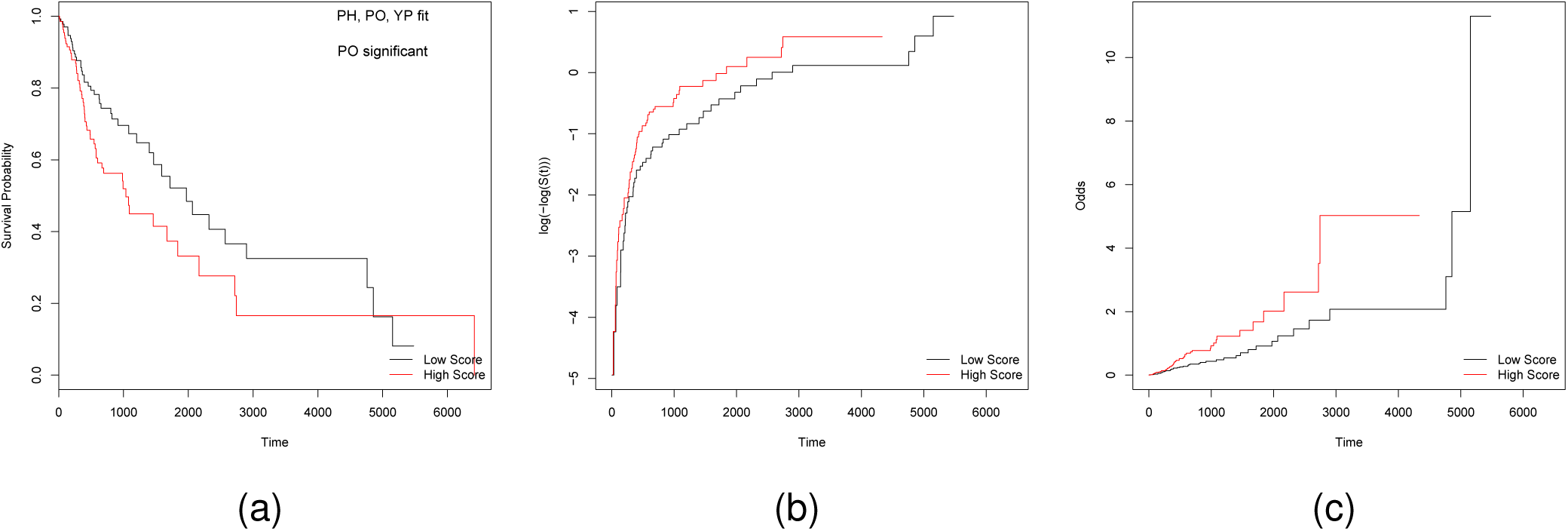
(a) Kaplan-Meier survival curves, (b) Cumulative hazard curves (log-scale), (c) Odds curves. Weighted expression of features selected by 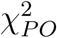 for data set 8

Next, we examine the differences between genomic features selected by various *R*^2^ measures, once again selecting the top 500 features in each case. SI: Figure 4 shows the overlaps between PO-based *R*^2^ measures - 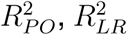, and 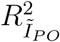 - for the glioblastoma data sets (1, 2 and 6). Since data set 2 contained a small number of features (454), the top 50 selected features were compared. The results are, however, consistent across these data sets where we observe a large fraction of common features selected by different measures. In all three data sets, a large fraction of features (> 80%) are common to all three measures as well as between any pair of measures. SI: Figure 5 shows the overlaps between PO-based *R*^2^ measures for the HNSCC (3, 7 and 8), ovarian (4) and oral (5) data sets where we, once again, observe a significant fraction of features common to different measures. Overall, 55-80% of features are common to all three measures while 61-96% of features are common between to any pair of measures. These results are not surprising since all three measures are based on the PO model and performed similarly in the simulations.

Venn diagrams corresponding to 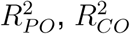, and 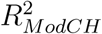 are shown in SI: Figures (data sets 1, 2 and 6) and 7 (data sets 3, 7, 8, 4 and 5). We observe that there are only minor overlaps between different measures across all data sets, with the largest overlaps occurring between 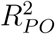 and 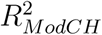 for data sets 5 and 8 and between 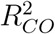 and 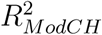 for data sets 1, 6 and 8. However, there are no features common to all three measures in data sets 3, 4, 6, 7 and 8, and only a very small number of common features in the remaining data sets. This is not surprising since each of these methods is based on a different model, so we would expect them to select different features. Although not shown here, we note that 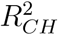 and 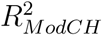 selected a large proportion of common features in all of the data sets; however, 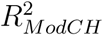 provided a robust estimate compared to 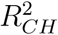 due to its computational correction. The differences in observed selected features across the three measures are most likely related to the presence of NPH features; for example, 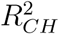 is specifically designed to identify crossing hazards, but a measure based on PO or YP will provide more flexibility as it allows for different types of time-varying hazards as well as PH. Thus, the appropriate measure can be chosen based on the goal of feature selection and type of hazards present or expected. We emphasize that our proposed measures provide a more versatile and general framework that allows for inclusion of various types of hazards.

The *R*^2^ measures can be interpreted as the percentage of separation in feature expression between those experiencing the event of interest and those not experiencing it. As seen in SI: Figures 6 and 7, 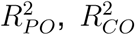, and 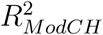 select fairly independent subsets of features, and each set can be used for further exploration and study. The PO model-based measures demonstrated their ability to handle PH and various forms of NPH throughout the simulation results while 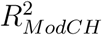 only performed well in detecting crossing hazards; moreover, 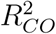 offers an alternative approach to 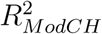 and is better suited for handling particular forms of time-varying hazards. Thus, each measure provides useful information specific to a particular model of interest while PO model-based *R*^2^ measures provide overall flexibility relative to other measures.

## 7 Summary and discussion

In this paper, we proposed unified methods for feature selection in large-scale genomic data in the presence of censored survival outcomes. We illustrated the utility of these methods using real-life studies in cancer genomics; in particular, we demonstrated their ability to handle the challenges due to various forms of non-proportional hazards. The proposed methods are based on models that relax the PH assumption and are able to identify genomic features with a time-varying effect with increased specificity and sensitivity.

Our methods are flexible and generalize existing methods for feature selection by casting them within unifying frameworks. First, we proposed a general framework to test for the effect of a genomic feature under the YP model using KL information divergence by developing the measure *I*_*Y P*_ which quantifies this effect. *I*_*Y P*_ contains corresponding measures for the PO and PH model - *I*_*PO*_ and *I*_*PH*_, respectively - as special cases. An advantage of these measures is that they do not require an estimate of the baseline hazard and, instead, are simple functions of model parameters. Using these measures, we developed a statistical test 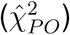 for genomic feature effect in the PO model where the test-statistic or *p*-value could be used for feature selection. Using *I*_*PH*_ and *I*_*PO*_, we developed 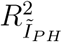 and 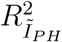 for the PH and PO model, respectively, that only rely on the corresponding regression coefficient; in addition, we developed alternative *R*^2^ measures 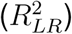 based on the likelihood ratio for the PO model. All these *R*^2^ measures are easily interpretable as the fraction of variation explained by the respective models. Finally, we proposed a generalized pseudo-*R*^2^ measure that embeds measures for the PO, CO, PH, and CH models as special cases. These measures do not require an estimate of the model parameter and can be easily interpreted as the percentage of separability between subjects in the event and non-event groups according to feature expression.

All proposed *R*^2^ measures can be applied to quantitative (continuous or ordered categorical) data and *I* measures are applicable to quantitative or dichotomized data; however, *I*_*Y P*_ is currently usable only on dichotomized data. Use of (appropriately variance stabilized and normalized) quantitative feature expression aids in robust estimation of time-varying effects and interpretability while dichotomized expression facilitates visualization of results. However, it is important to be aware of the potential effect of dichotomization - using the median split or any other arbitrary cut-off - on the PH assumption. This is evidenced by the results in Tables 1 and 2 for data sets 1 and 2 where dichotomization has a negative and positive effect, respectively, on GOF. The proposed *I* measures are useful for feature ranking in general and, in particular, when the distribution of feature expression or a normalizing transformation for it is unknown. Furthermore, *I*_*PO*_, 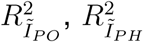 and 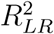 can accommodate potential confounders such as age and stage of disease directly or indirectly in their computation (as illustrated in our examples) as well as a group of pre-selected features depending on the application of interest. While *I*_*Y P*_ offers an approach for feature ranking using a flexible survival model for dichotomized feature expression, 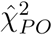 provides a method for feature ranking as well as selection for both continuous and dichotomized feature expression. Typically, it selects more features at the same significance threshold and, thus, provides a more lenient approach for feature discovery relative to standard methods and CON as evidenced in Table 4. Moreover, our results demonstrate that it includes a significant fraction of features identified by CON. This is a desirable property of any feature selection method and it enables appropriate correction for multiple testing through use of FDR based not only on the Benjamini-Hochberg approach which assumes independent hypotheses but even a more stringent method such as Benjamini-Yekutieli that accounts for dependence amongst hypotheses. Such flexibility is not possible with currently available methods for feature selection and ranking especially within a broad framework that includes the YP, PO and PH models.

Our extensive simulation studies demonstrated that there were a variety of scenarios where our proposed measures outperformed currently available methods. An important consideration is that when marginal screening methods are utilized for the purpose of feature ranking and selection, the issue of multiple testing becomes less important in comparison to adjusting for potential confounders when considering different models and measures. The proposed methods demonstrated their ability to correctly select genomic features associated with survival in the presence of different types of time-varying effects in real genomic data, after adjusting for potential confounders and for multiple testing, as well as in simulated data. As genomic technologies continue to advance and as more clinical, demographic and genomic data are generated and stored in repositories such as TCGA, Gene Expression Omnibus (https://www.ncbi.nlm.nih.gov/geo/) etc., feature selection methods will become increasingly important as we attempt to identify genomic features with a prognostic impact on patient survival. Although we focused on genomic feature selection in this paper, it should be noted that the proposed methods are directly applicable to a broad array of high-throughput “omics” studies such as those involving genome-wide association, proteomics, metabolomics, transcriptomics and radiomics. In particular, radiomics is a rapidly developing area which involves a multitude of quantitative measurements of tumor heterogeneity based on various imaging modalities such as computed tomography and magnetic resonance imaging. There has been considerable interest recently in correlating intra-tumor heterogeneity based on textural features to survival endpoints.

## Acknowledgement

The work of KD was funded in part through the NIH/NCI Cancer Center Support Grant P30 CA006927. An earlier version of this work won the student paper award (LS) in the Statistical Learning and Data Science Section of the Joint Statistical Meetings in Baltimore, 2017.

## Supplementary Information

### 8 Data Sets and Implementation

#### Data sets 1, 2 and 6

These data sets were published by TCGA (http://cancergenome.nih.gov/). Data set 1 consists of the methylation profiles (beta values) of tumor samples from 280 patients with glioblastoma (GBM) obtained using the Infinium HumanMethylation27 platform. The beta values were normalized using the logit transformation. For genes with multiple methylation probes, the probe most negatively correlated with expression is used. Data set 2 consists of microRNA expression profiles in the form of z-scores compared to all tumors for 416 patients with GBM. Data set 6 consists of merged mRNA and microRNA z-scores from 426 patients with GBM where mRNA expression z-scores were compared to diploid tumors (diploid for each gene) using median values from all three mRNA expression platforms (Affymetrix U133, Affymetrix Exon, and Agilent), and microRNA z-scores were compared to all tumors. Data sets 1, 2 and 6 contain a total of 9,452, 454 and 15,546 features, respectively.

#### Data sets 3, 7 and 8

These data sets were published by TCGA (http://cancergenome.nih.gov/). The raw genome-wide methylation data for ∼485, 000 CpG sites (based on Infinium HumanMethylation450 BeadChip Kit, Illumina, Inc.) obtained from tumor samples of 286 patients with head and neck squamous cell carcinoma (HNSCC) were retrieved from TCGA, and M-values (methylation signal quantified by logit-transformed beta values) were calculated for each CpG site using the Bioconductor package Minfi (Aryee et al., 2014). CpGs located in the transcription start sites and UTR regions for each gene were retrieved and used in further analyses. Somatic copy number variation (CNV) - expressed in discretized form as gain or loss (−2,-1,0,1,2) - and RNA-Seq gene expression data - presented as RSEM values (Li & Dewey, 2011) - were obtained from the Broad Institute (http://gdac.broadinstitute.org/). CNV data was filtered by removing genes with identical copy number variation across subjects. RNA-seq data was normalized using the log_2_(*x* + 1) transformation. A gene was included in the analyses only if (i) if 50% of patients have expression values for that gene, and (ii) protein expression of that gene was observed in at least one head and neck cancer sample in the Human Protein Atlas database (Uhlen et al., 2015). Data sets 7 and 8 consist of methylation and CNV data, respectively, for 286 patients with HNSCC while data set 3 consists of RNA-Seq data for 221 patients with cancers of the oral cavity, a subgroup of HNSCC. Data sets 3, 7 and 8 contain a total of 19,341, 49,270 and 5,869 features, respectively, after the above pre-processing steps.

#### Data sets 4

Tothill et al. (2008) studied the relationship between recurrence-free survival and gene expression in ovarian cancer using tumor samples from 276 subjects and Affymetrix U133 Plus 2 microarrays. This RMA normalized data set (Irizarry et al., 2003) was filtered using a coefficient of variation threshold of 35% to remove genes with low variation in expression and contains the expression profiles of 24,739 probe sets. In all analyses, log_2_ transformed data was used.

#### Data set 5

Saintigny et al. (2011) studied 86 subjects enrolled in a clinical chemoprevention trial where the primary endpoint of interest was the development of oral cancer. This RMA normalized and log_2_ transformed data set (Irizarry et al., 2003) contains the expression profiles of 12,776 probe sets obtained using the Human Gene 1.ST platform.

#### Implementation

All computations were done using the R Statistical Language and Environment (R Core Team, 2018) and Bioconductor (Gentleman et al., 2004). The following packages were utilized as needed: survival, timereg, YPmodel, qvalue, Minfi, affy, concreg, gplots, VennDiagram and latex2exp.

### 9 Methods

#### 9.1 Derivation of *I*_*Y P*_

Under the YP model defined in equations (2.5) and (2.6) and the identities *f* (*t*|*z*) = *λ*(*t*|*z*)*S*(*t*|*z*) and *f*_0_(*t*) = *λ*_0_(*t*)*S*_0_(*t*), feature-specific KL information divergence for this model has the form

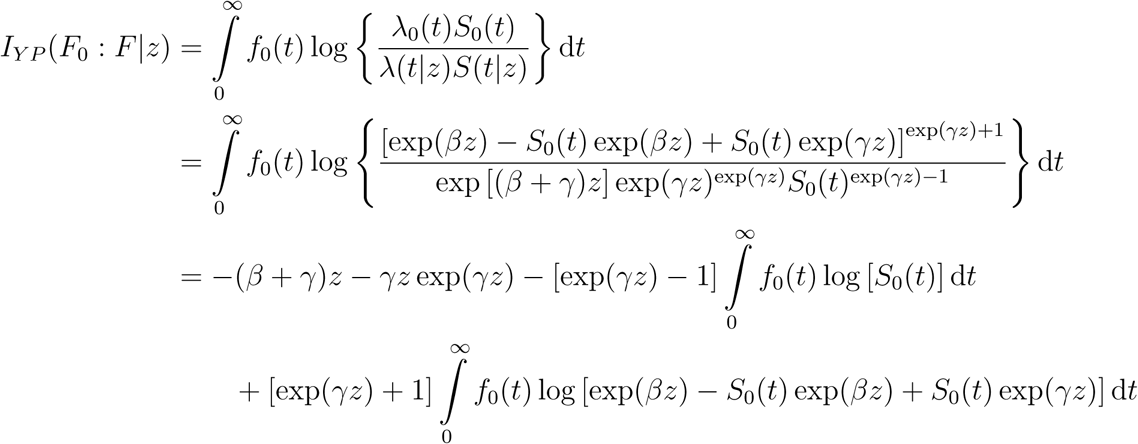

In the second term, let *u* = *S*_0_(*t*) and d*u* = −*f*_0_(*t*)d*t.*

In the third term, let *u* = exp(*βz*) − *S*_0_(*t*) exp(*βz*) + *S*_0_(*t*) exp(*γz*)

and d*u* = *f*_0_(*t*) [exp(*βz*) − exp(*γz*)] d*t*

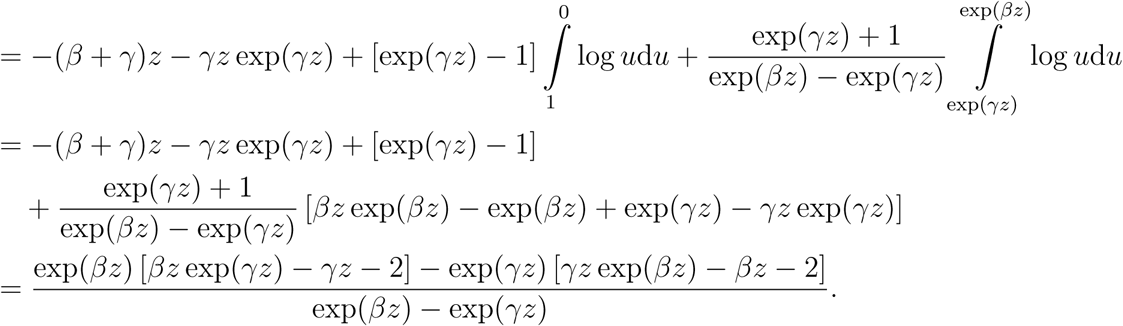

##### Theorem 1.

*Î*_*Y P*_ *is a maximum likelihood estimator and is asymptotically normal with mean I*_*Y P*_.

*Proof.* It can be seen from equation (4.3) that *Î*_*Y P*_ is a simple transformation of the pseudo maximum likelihood estimators 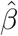 and 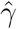. Since 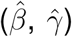 is asymptotically bivariate normal with mean (*β, γ*), using the invariance property of maximum likelihood estimators it can be concluded that *Î*_*Y P*_ is also asymptotically normal with the mean above. For more details, see Yang & Prentice (2005) and Devarajan & Ebrahimi (2009).

□

#### 9.2 Derivation of *I*_*PO*_ and *V ar*(*Î*_*PO*_)

*I*_*PO*_(*F*_0_ : *F*) is obtained by setting *γ* = 0 in the expression for *I*_*Y P*_ (*F*_0_ : *F*). It turns out that KL information divergence is symmetric for the PO model, i.e., *I*_*PO*_(*F*_0_ : *F*) = *I*_*PO*_(*F* : *F*_0_) (Spirko, 2016). This measure is weighted equally towards the null and alternative hypotheses.

##### Theorem 2.

*Î*_*P O*_ *is a maximum likelihood estimator and is asymptotically normal with mean I*_*PO*_.

*Proof.* It can be seen from equation (4.4) that *Î*_*PO*_ is a simple transformation of the modified partial likelihood estimator 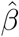. Since 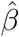 is asymptotically normal with mean *β*, using the invariance property of maximum likelihood estimators it can be concluded that *Î*_*PO*_ is also asymptotically normal with the mean above. For more details, see Martinussen & Scheike (2006) and Devarajan & Ebrahimi (2009).

□

Using equation (4.4), for a given feature *j* with expression *z*, variance of *Î*_*PO*_ can be written in terms of 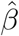 as (suppressing the subscript *j*)

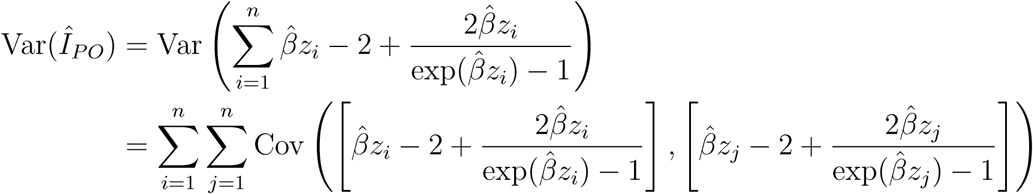

Let *I*(*β*) denote *I*_*PO*_ in equation (4.4) expressed as a function of the parameter *β* for each observation *i*. Expanding *I*(*β*) using the first three terms of the Taylor series, we get

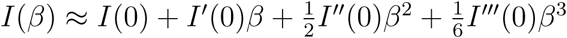

where

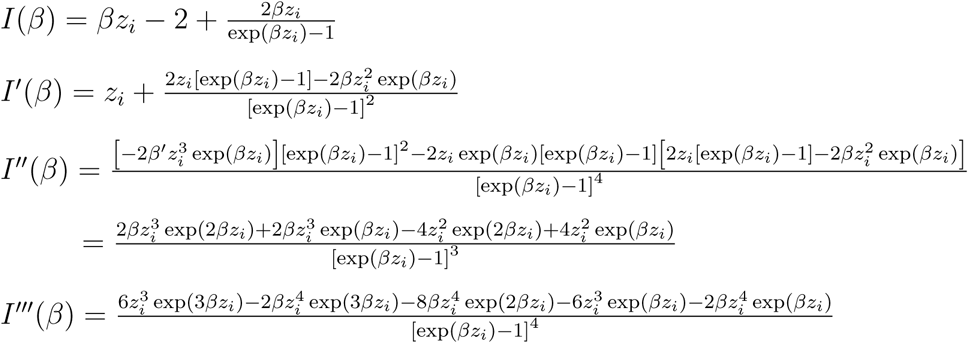

Taking the limit as *β* → 0 of all four functions above, we get

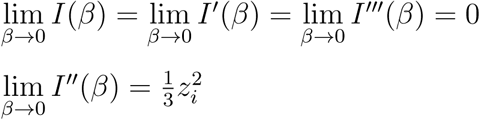

Thus, 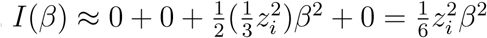

Now, using the above results the approximate variance of *Î*_*PO*_ can be computed as follows.

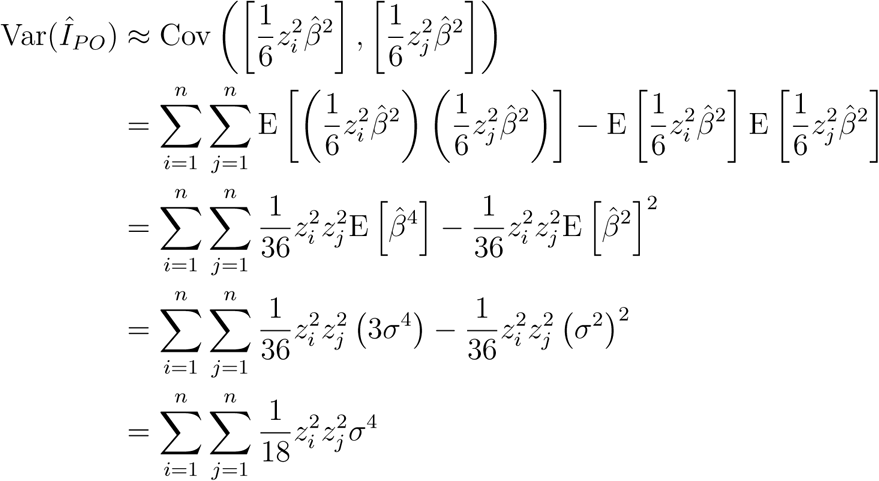

The fourth equality above is obtained using the fact 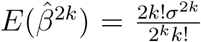 where *σ*^2^ is the variance of 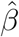 under *H*_0_.

#### 9.3 Derivation of *Ĩ*_*PO*_ and *Ĩ*_*PH*_

##### PO model

For the PO model, using equations (4.9) and (4.7),

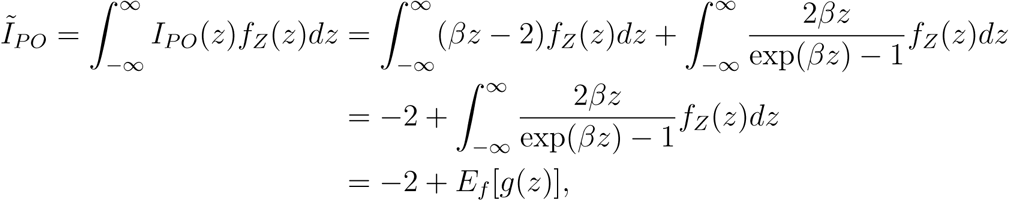

where 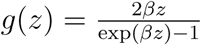. Using the Taylor series expansion, we can estimate *E*_*f*_ [*g*(*z*)] by

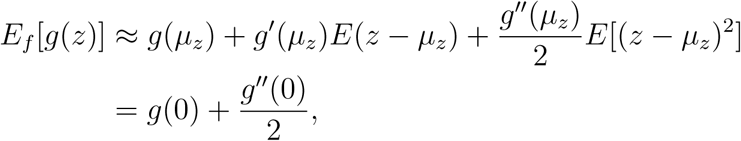

where 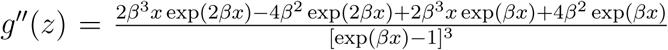. Now, take the 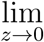 for *g*(*z*) and *g*′′(*z*) using L’Hopital’s rule to get

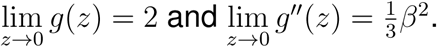

Thus, 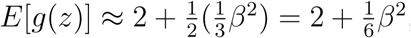, and therefore,

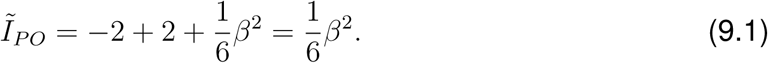

The *R*^2^ measure is then defined as

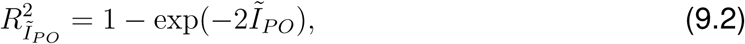

where *β* in *Ĩ*_*PO*_ is replaced by 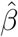, the modified partial likelihood estimator in the PO model (Martinussen and Scheike 2006).

##### PH model

For the PH model, using equations (4.8) and (4.9),

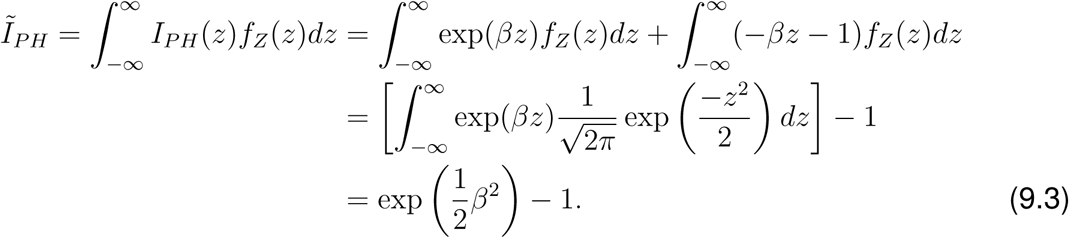

*R*^2^ is defined as

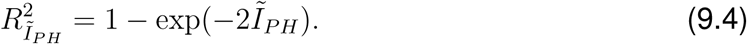

where *β* in *Ĩ*_*PH*_ is replaced by 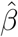, the partial likelihood estimator parameter in the PH model (Cox 1972).

#### 9.4 Derivation of the generalized pseudo-*R*^2^ measure

Following standard counting process notation for censored survival data, let *N*_*i*_(*t*) = {0, 1} denote the number of events that have occurred for subject *i, i* = 1,, *n* in the interval (0, *t*] and let *Y*_*i*_(*t*) = 1 indicate that subject *i* is at risk just before time *t*. Here, 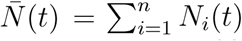 and 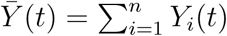 denote, respectively, the total number of events that have occurred in (0, *t*] and the number of subjects at risk at time *t*. Let *β* denote the parameter and let *U*_*i*_(*β*; *t*) denote the score function for individual *i* obtained as the derivative of the logarithm of partial likelihood with respect to *β* for a particular model of interest. For each model, it can be shown that the score evaluated at *β* = 0 has the form

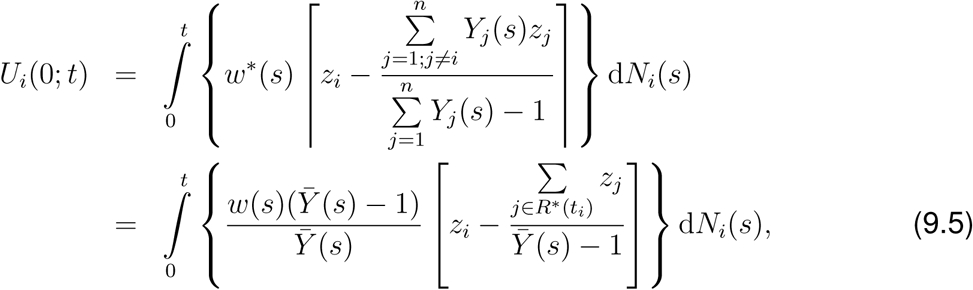

where 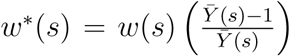, *w*(*s*) is a weight function that is model-specific, and *R*^*^(*t*) is the set of individuals not experiencing the event at time *t*. Thus, it can be seen that this is a measure of the weighted difference in the expression of a genomic feature between subjects observed to experience the event of interest and those observed to not experience the event. For each observation *i*, an estimate for *U*_*i*_(0; *t*) is given by

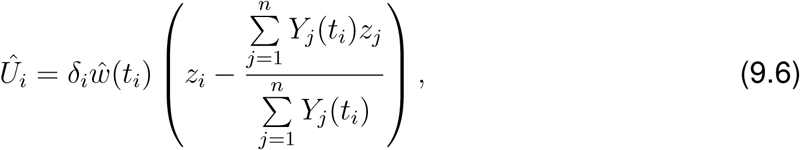

where estimation of *ŵ*(*t*_*i*_) is discussed later in this section and *δ*_*i*_ is the indicator of failure at time *t*_*i*_. Following Rouam et al. (2011), we utilize the robust score proposed in Lin & Wei (1989) given by

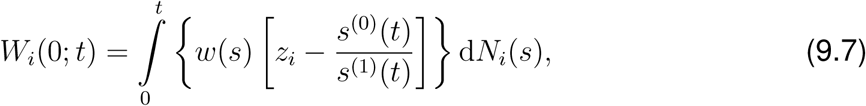

where 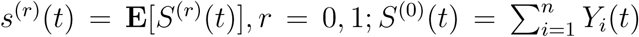 and 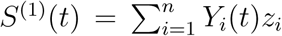. This quantity can be estimated by

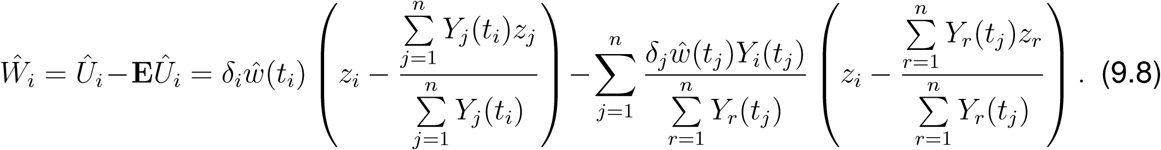

The sum of *W*_*i*_s is identical to the sum of *U*_*i*_s but *W*_*i*_s are independent under the PH model. Using *W*_*i*_, a pseudo-*R*^2^ index can then be written as

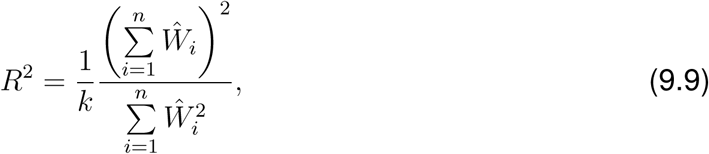

where *k* is the number of uncensored failure times. Next, we compute *U* (0; *t*) for the PO and CO models defined in equations (2.3) and (2.4), respectively, and use it to derive respective weights *w*(*t*).

##### PO model

For the PO model defined in equation (2.3), the following quantities represent the survival and hazard functions, respectively,

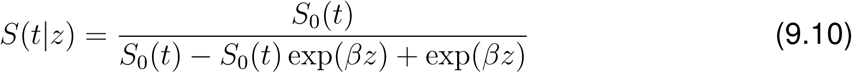

and

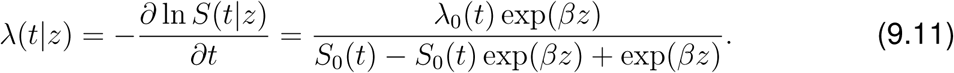

Using equations (9.10) and (9.11), the partial likelihood for the PO model can be written as

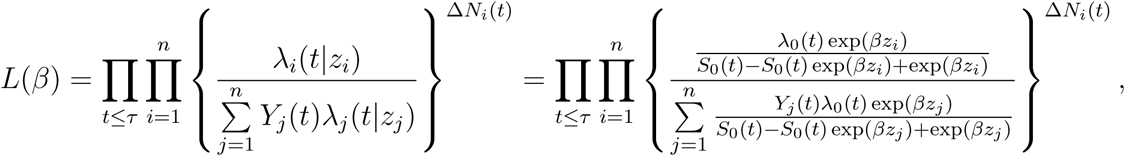

where *Y*_*j*_(*t*) = 1 if the subject is at risk before time *t*, and *z*_*i*_ represents the expression of a given feature for subject *i*. For fixed *t*, 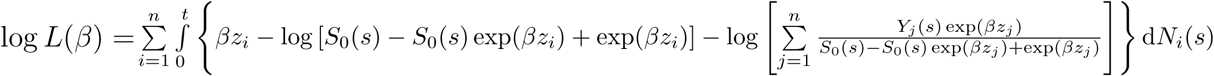

Hence,

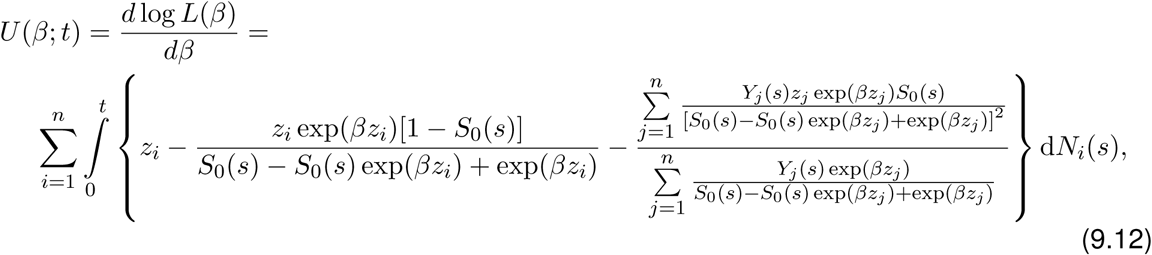

and setting *β* = 0, we get

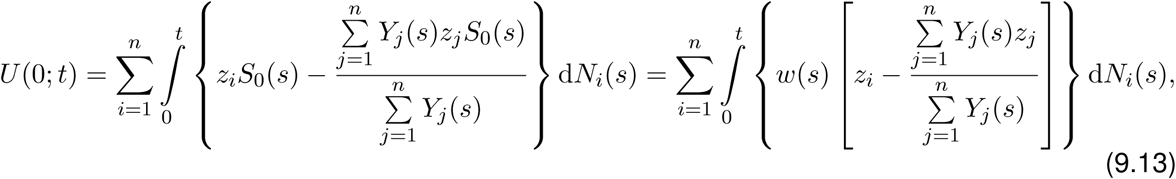

where *w*(*s*) = *S*_0_(*s*).

##### CO model

The CO model in equation (2.4) introduces another parameter into the PO model and has the form

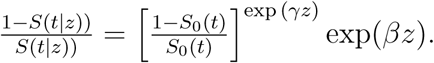

It is useful because of this generalization, and it allows the hazard functions corresponding to two values of a covariate to cross. Here, we set *γ* = *β* such that

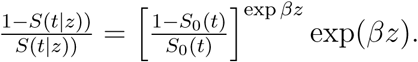

Then, the survival and hazard functions for this special case of the CO model can be written as

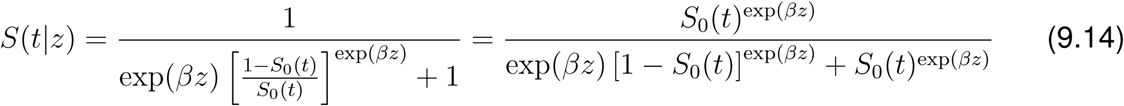

and

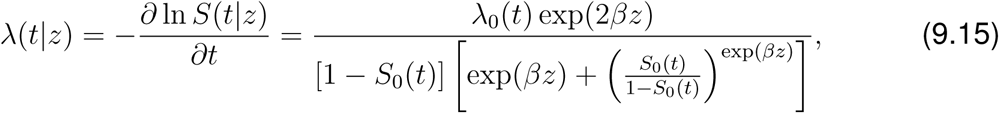

respectively. The partial likelihood is written as

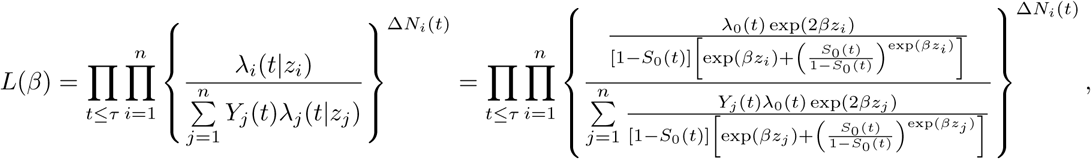

where *Y*_*j*_(*t*) = 1 if the subject is at risk before time *t*. For fixed *t*, log *L*(*β*) =

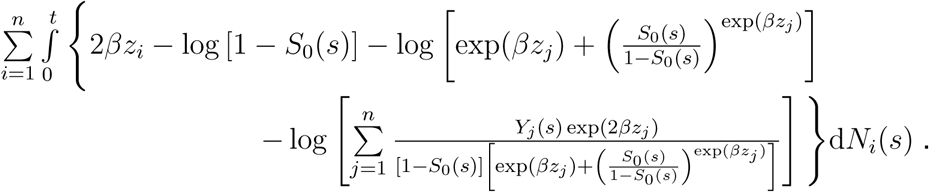

Hence,

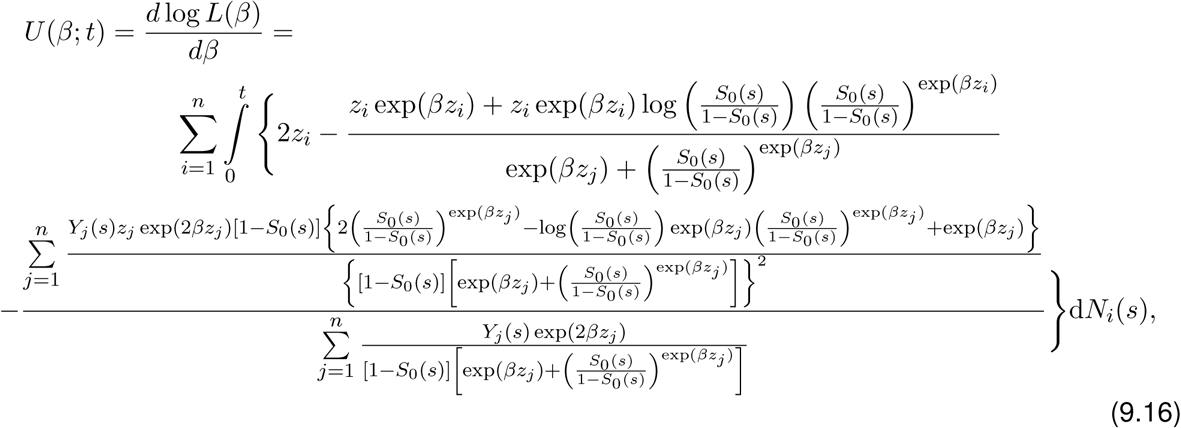

and setting *β* = 0, we get

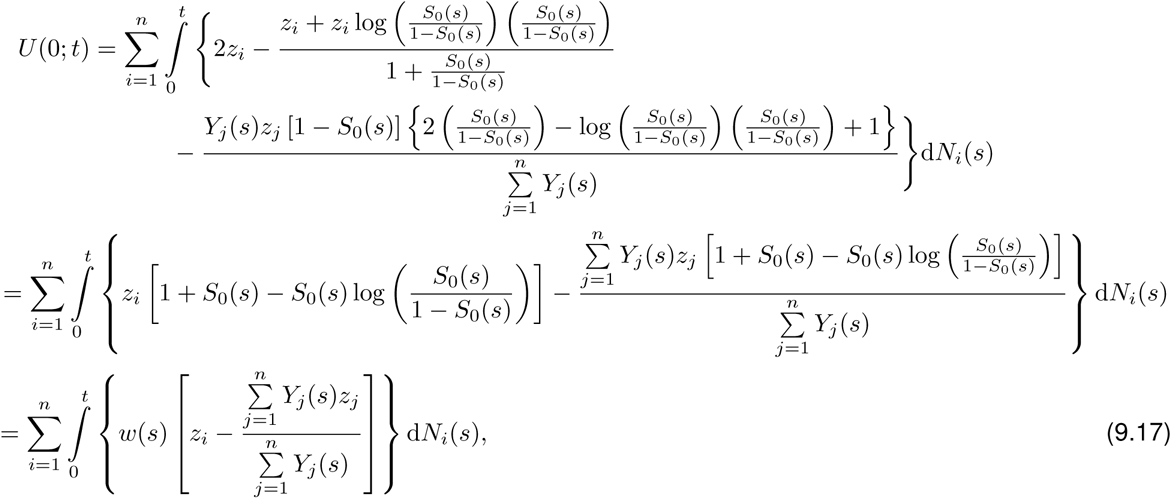

where 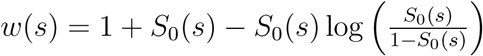.

##### General Form

Using equations (9.13) and (9.17), the score function can be expressed in the following generalized form

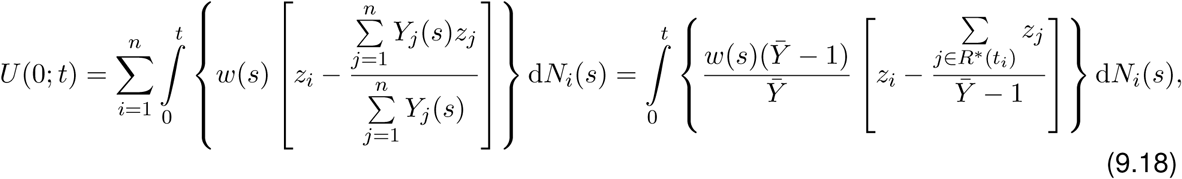

that includes the PO and CO models where *w*(*s*) is the model-specific weight function and the other terms are as defined before. Using weights specified in Table 3 for the PH and CH models (Rouam et al, 2010; 2011), equation (9.18) can be seen to represent the generalized score function that includes the PH, CH, PO and CO models under consideration in this paper.

##### Computational correction for 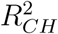 and 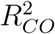

The measure 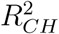 has weight 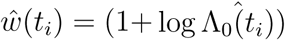, where Λ_0_ (*t*_*i*_) is estimated by the left-continuous version of Nelson’s estimator. However, this weight has an inherent numerical issue when 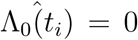. To handle this error, Rouam et al. (2011) set *ŵ*(*t*_*i*_) = 1 if 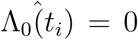, implying that 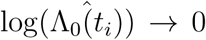 as 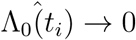. This is unrealistic because 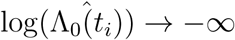 as 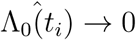. Thus, we propose an empirical correction for this error that uses a plot of the cumulative hazard versus the weight to obtain an approximation for the weight as the cumulative hazard approaches zero. In our computations, we set *ŵ*(*t*_*i*_) equal to this approximation when 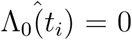. We call this modified measure 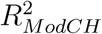. Similarly, for 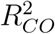, an empirical correction was made to account for the numerical issue when 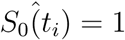 by obtaining a graphical approximation for the weight as 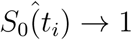.

### 10 Simulation Studies

#### 10.1 Simulation scheme 1

First, we look at the results for scheme 1, the univariate approach. Table 5 reports the AUCs for each method across the five models considered - LN, LL1, LL2, W1 and W2. We note that the standard deviations of the AUCs were uniformly very small throughout and ranged from 7*X*10^−4^ to 0.02. In Table 5, we observe several scenarios where the proposed measures outperform existing methods.

First, we look at the performance of the three *R*^2^ measures based on the PO model - 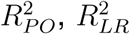, and 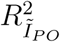. In each scenario and across all censoring schemes, these measures perform almost identically. In some instances, such as LL2 under 0% and 33% censoring, we do observe a slight improvement in 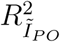. Next, we consider PH model-based measures - the newly proposed 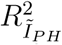 and the existing 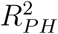. From Table 5, we observe that the AUCs are almost identical for these measures, with a slight improvement noted in some cases for 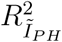. In fact, based on the Youden indices in Table 7, we note that the proposed 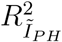 outperforms 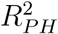 for all three censoring proportions for LL2 and for the 0% censoring case for LN; this also holds true for AUCs.

**Table 6:** Simulation scheme 2: Comparison of methods (AUC)

| Censoring | Measure | LN | LL1 | LL2 | W1 | W2 |
| --- | --- | --- | --- | --- | --- | --- |
| 0% | $I_{PO}$ | 0.84 | 0.91 | 0.54 | 0.91 | 0.90 |
| | $I_{YP}$ | 0.86 | 0.89 | 0.77 | 0.89 | 0.89 |
| | $\hat{c}'_+$ | 0.82 | 0.91 | 0.50 | 0.91 | 0.90 |
| | $R_{PO}^2$ | 0.84 | 0.91 | 0.55 | 0.91 | 0.90 |
| | $R_{LR}^2$ | 0.84 | 0.91 | 0.56 | 0.91 | 0.90 |
| | $R_{I_{PO}}^2$ | 0.82 | 0.91 | 0.57 | 0.91 | 0.90 |
| | $R_{CO}^2$ | 0.76 | 0.51 | 0.88 | 0.51 | 0.88 |
| | $R_{CH}^2$ | 0.89 | 0.86 | 0.89 | 0.86 | 0.79 |
| | $R_{ModCH}^2$ | 0.89 | 0.87 | 0.89 | 0.86 | 0.81 |
| | $R_{I_{PH}}^2$ | 0.55 | 0.91 | 0.63 | 0.91 | 0.90 |
| | $R_{PH}^2$ | 0.55 | 0.90 | 0.66 | 0.91 | 0.89 |
| 33% | $I_{PO}$ | 0.87 | 0.91 | 0.62 | 0.91 | 0.90 |
| | $I_{YP}$ | 0.84 | 0.85 | 0.82 | 0.88 | 0.86 |
| | $\hat{c}'_+$ | 0.80 | 0.91 | 0.50 | 0.91 | 0.90 |
| | $R_{PO}^2$ | 0.87 | 0.90 | 0.63 | 0.90 | 0.89 |
| | $R_{LR}^2$ | 0.88 | 0.91 | 0.61 | 0.91 | 0.90 |
| | $R_{I_{PO}}^2$ | 0.86 | 0.91 | 0.64 | 0.91 | 0.90 |
| | $R_{CO}^2$ | 0.59 | 0.51 | 0.87 | 0.53 | 0.88 |
| | $R_{CH}^2$ | 0.89 | 0.87 | 0.90 | 0.87 | 0.70 |
| | $R_{ModCH}^2$ | 0.88 | 0.87 | 0.89 | 0.87 | 0.74 |
| | $R_{I_{PH}}^2$ | 0.72 | 0.90 | 0.57 | 0.90 | 0.90 |
| | $R_{PH}^2$ | 0.75 | 0.90 | 0.54 | 0.90 | 0.89 |
| 67% | $I_{PO}$ | 0.89 | 0.90 | 0.80 | 0.90 | 0.86 |
| | $I_{YP}$ | 0.69 | 0.65 | 0.68 | 0.69 | 0.68 |
| | $\hat{c}'_+$ | 0.84 | 0.90 | 0.53 | 0.90 | 0.87 |
| | $R_{PO}^2$ | 0.89 | 0.90 | 0.81 | 0.90 | 0.84 |
| | $R_{LR}^2$ | 0.89 | 0.90 | 0.80 | 0.90 | 0.89 |
| | $R_{I_{PO}}^2$ | 0.88 | 0.90 | 0.80 | 0.90 | 0.89 |
| | $R_{CO}^2$ | 0.56 | 0.50 | 0.71 | 0.50 | 0.87 |
| | $R_{CH}^2$ | 0.81 | 0.88 | 0.88 | 0.88 | 0.53 |
| | $R_{ModCH}^2$ | 0.81 | 0.88 | 0.87 | 0.88 | 0.55 |
| | $R_{I_{PH}}^2$ | 0.87 | 0.90 | 0.70 | 0.90 | 0.89 |
| | $R_{PH}^2$ | 0.89 | 0.90 | 0.71 | 0.90 | 0.87 |

**Table 7:** Simulation schemes 1 & 2: Comparison of methods (Youden Index)

| Scheme | 1 |  |  |  |  |  | 2 |  |  |  |  |  |
| --- | --- | --- | --- | --- | --- | --- | --- | --- | --- | --- | --- | --- |
| Censoring | 0% |  | 33% |  | 67% |  | 0% |  | 33% |  | 67% |  |
| Measure | LN | LL2 | LN | LL2 | LN | LL2 | LN | LL2 | LN | LL2 | LN | LL2 |
| $I_{PO}$ | 0.53 | 0.34 | 0.61 | 0.42 | 0.68 | 0.50 | 0.23 | 0.03 | 0.24 | 0.09 | 0.26 | 0.24 |
| $I_{YP}$ | 0.99 | 0.49 | 0.49 | 0.22 | 0.38 | 0.00 | 0.22 | 0.17 | 0.12 | 0.16 | 0.07 | 0.04 |
| $\hat{c}'_+$ | 0.53 | 0.33 | 0.54 | 0.28 | 0.61 | 0.29 | 0.21 | 0.02 | 0.19 | 0.00 | 0.22 | 0.01 |
| $R^2_{PO}$ | 0.53 | 0.31 | 0.62 | 0.39 | 0.68 | 0.48 | 0.23 | 0.04 | 0.25 | 0.10 | 0.26 | 0.24 |
| $R^2_{LR}$ | 0.53 | 0.33 | 0.60 | 0.41 | 0.68 | 0.49 | 0.23 | 0.05 | 0.25 | 0.07 | 0.26 | 0.22 |
| $R^2_{I_{PO}}$ | 0.52 | 0.36 | 0.59 | 0.43 | 0.68 | 0.50 | 0.20 | 0.06 | 0.23 | 0.11 | 0.25 | 0.19 |
| $R^2_{CO}$ | 0.89 | 0.53 | 0.41 | 0.43 | 0.24 | 0.26 | 0.17 | 0.23 | 0.06 | 0.22 | 0.07 | 0.14 |
| $R^2_{CH}$ | 0.99 | 0.59 | 0.82 | 0.56 | 0.60 | 0.52 | 0.26 | 0.28 | 0.25 | 0.28 | 0.23 | 0.28 |
| $R^2_{ModCH}$ | 0.99 | 0.61 | 0.80 | 0.58 | 0.60 | 0.54 | 0.26 | 0.28 | 0.24 | 0.28 | 0.23 | 0.28 |
| $R^2_{I_{PH}}$ | 0.66 | 0.04 | 0.49 | 0.18 | 0.66 | 0.44 | 0.02 | 0.06 | 0.13 | 0.03 | 0.24 | 0.13 |
| $R^2_{PH}$ | 0.46 | 0 | 0.49 | 0.12 | 0.66 | 0.41 | 0.04 | 0.09 | 0.17 | 0.02 | 0.25 | 0.16 |

Next, we examine the LN case where we observe from Table 5 that 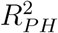 and 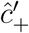are outperformed by various measures under different scenarios. Specifically, 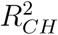 and 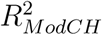 perform similarly and outperform PO and PH model-based *R*^2^ measures, except for the 67% censoring case where 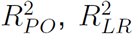, and 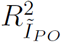 perform the best. This is not surprising since the LN model allows for crossing hazards. *R*^2^ performs well for lower censoring proportions, but its AUC decreases as censoring increases. As censoring increases, PO model-based *R*^2^ measures also outperform PH model-based measures in terms of AUCs as well as Youden indices shown in Table 7. *I*_*Y P*_ outperforms *I*_*PO*_ and 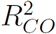 when the censoring proportion is 0 and 33% but its performance decreases as censoring increases. It should be noted that *I*_*Y P*_ can only be applied to dichotomized genomic data; thus, *I*_*Y P*_ ‘s performance may be affected by its inability to accommodate continuous data. *I*_*PO*_ also outperforms 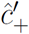 as censoring increases which is also evidenced by the Youden indices in Table 7.

Under LL1, we expect PO model-based measures to perform well since the log-logistic model is related to the PO model, and our results provide evidence in support. While 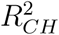 and 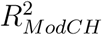 do improve as censoring increases, 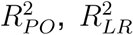, and 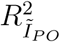 still outperform 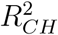 and 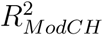 at each censoring level. Similar to the LN case, the performance of 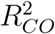 and *I*_*Y P*_ decreases as censoring increases. Under LL2, 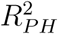 and 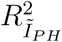 perform significantly worse than other *R*^2^ measures, but their AUCs do increase from approximately .50 − .54 in the 0% censoring case to .78 in the 67% censoring case. Thus, while these measures show improved performance with higher censoring, they are still consistently outperformed by other model-based measures. This is also evident from Table 7, where 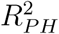 and 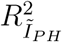 have the lowest Youden index of all the reported measures with the exception of 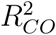 under 67% censoring. The log-logistic model of LL2 allows for crossing hazards and is related to the CO model and, not surprisingly, 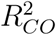 outperforms PO model-based *R*^2^ measures in the 0 and 33% censoring cases but, similar to the LN and LL1 models, we see its AUCs drop for 67% censoring. 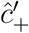 is outperformed by *I*_*PO*_ and/or *I*_*Y P*_, as well as by 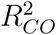, at each censoring level, with *I*_*Y P*_ performing better for lower censoring proportion and *I*_*PO*_ performing better for higher censoring proportion. In this case, we also observe that PO model-based *R*^2^ measures outperform PH model-based methods which emphasizes the PO model’s ability to handle some forms of NPH. These observations are further supported by the Youden indices shown in Table 7.

Under the W1 model, we observe that 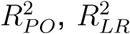, and 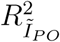 outperform 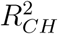 and 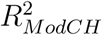 at all censoring levels. This result is intuitive because this Weibull model is related to the PH model and the PO model does allow for PH. Under the W2 model, we observe that 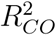outperforms 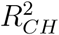 and 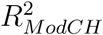 at all censoring levels, especially for higher censoring proportions where the improvement in performance is markedly higher. Furthermore, PO model-based measures outperform 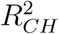. The W2 model allows for crossing hazards, and yet here, we observe a clear advantage for our PO and CO model-based measures over 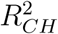 which was purposefully designed to handle crossing hazards. Also, in this case we observe that 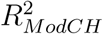, the proposed modification to 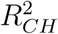, performs significantly better than 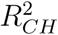 as the censoring proportion increases. Youden indices for models W1, W2, and LL1 under scheme 1 are listed in Table 8 and support the results in Table 5.

**Table 8:** Simulation Scheme 1: Comparison of methods (Youden Index)

| Censoring | 0% |  |  | 33% |  |  | 67% |  |  |
| --- | --- | --- | --- | --- | --- | --- | --- | --- | --- |
| Measure | LL1 | W1 | W2 | LL1 | W1 | W2 | LL1 | W1 | W2 |
| $I_{PO}$ | 0.92 | 0.95 | 0.98 | 0.88 | 0.92 | 0.92 | 0.80 | 0.85 | 0.72 |
| $I_{YP}$ | 0.73 | 0.92 | 0.98 | 0.41 | 0.73 | 0.86 | 0.16 | 0.34 | 0.70 |
| $\hat{c}'_+$ | 0.92 | 0.95 | 0.97 | 0.87 | 0.93 | 0.94 | 0.76 | 0.84 | 0.81 |
| $R^2_{PO}$ | 0.92 | 0.94 | 0.97 | 0.87 | 0.92 | 0.91 | 0.80 | 0.85 | 0.72 |
| $R^2_{LR}$ | 0.92 | 0.98 | 0.97 | 0.88 | 0.95 | 0.92 | 0.81 | 0.86 | 0.73 |
| $R^2_{\tilde{I}_{PO}}$ | 0.92 | 0.98 | 0.98 | 0.88 | 0.95 | 0.92 | 0.81 | 0.86 | 0.73 |
| $R^2_{CO}$ | 0.48 | 0.63 | 0.99 | 0.22 | 0.48 | 0.61 | 0.15 | 0.05 | 0.64 |
| $R^2_{CH}$ | 0.19 | 0.14 | 0.80 | 0.49 | 0.25 | 0.32 | 0.63 | 0.58 | 0.10 |
| $R^2_{ModCH}$ | 0.27 | 0.22 | 0.80 | 0.52 | 0.31 | 0.44 | 0.63 | 0.59 | 0.30 |
| $R^2_{\tilde{I}_{PH}}$ | 0.88 | 0.99 | 1.00 | 0.87 | 0.97 | 0.98 | 0.81 | 0.87 | 0.80 |
| $R^2_{PH}$ | 0.87 | 0.96 | 1.00 | 0.85 | 0.94 | 0.97 | 0.79 | 0.86 | 0.79 |

#### 10.2 Simulation scheme 2

Next, we consider results from simulation scheme 2 which are shown in Table 6. In general, AUCs are observed to be slightly lower than those in Table 5, but this is likely due to the complexity of the scheme itself where correlations are introduced between features. The observed trend in these results, however, mimic what was observed for scheme 1. We note that the standard deviations of AUCs were very small for this scheme as well, ranging from 7*X*10^−4^ to 0.02.

Similar to scheme 1, PO model-based *R*^2^ measures - 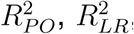, and 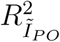 - perform almost identically across all scenarios and censoring levels. The PH model-based measures - 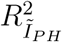 and 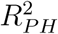 - also perform similarly. Under the LN model which allows for crossing hazards, 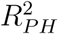 and 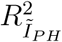 are outperformed by the proposed PO model-based *R*^2^ measures as well as by 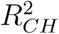 and 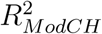 for 0 and 33% censoring cases, but for 67% censoring the PO model-based measures perform similarly and better than the other *R*^2^ measures. The AUCs for 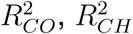 and 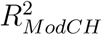 decrease as censoring increases, with PO model-based measures having a clear advantage for all censoring proportions. 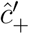 is also outperformed by *I*_*PO*_ and *I*_*Y P*_ for 0% and 33% censoring and by *I*_*P O*_ for 67% censoring. These differences are supported by the Youden indices shown in Table 7. Under the LL1 model, PO and PH model-based *R*^2^ measures perform similarly and result in slightly higher AUCs than 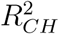 and 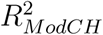. This result is also intuitive since the log-logistic model is related to the PO model which allows for both proportional hazards as well as some forms of non-proportional hazards. They also significantly outperform 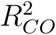 at each censoring level. *I*_*PO*_ and 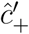 perform similarly and outperform *I*_*Y P*_, whose AUC decreases as censoring increases. Under the LL2 model, which allows for crossing hazards, 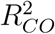 performs similarly to 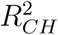 and 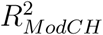 and they all significantly outperform PO and PH model-based *R*^2^ measures. This result is expected since the LL2 model allows for crossing hazards. However, similar to scheme 1, 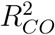 ‘s performance falls in the 67% censoring case. We also observe that the performance of 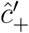 is poor and no better than a coin toss across all censoring proportions. *I*_*Y P*_ outperforms *I*_*PO*_ in the 0 and 33% censoring cases, but consistent with previous results, its AUC drops in the 67% censoring case. This is further corroborated by the Youden indices shown in Table 7 where *I*_*PO*_ is the best performer under 67% censoring. However, as alluded to earlier its performance may be affected by its inability to accommodate continuous genomic data. Under the W1 model, we observe results similar to that of scheme 1. 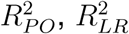, and 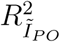 outperform 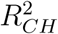 and 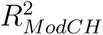 at all censoring levels which is intuitive since the PO model can accommodate proportional hazards and the Weibull model is related to the PH model. Under W2, 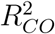 once again outperforms 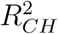 and 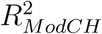 at all censoring levels and particularly for higher censoring where the improvement in performance is marked; in this case, we also observe that 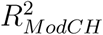 performs slightly better than 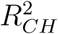 across all censoring proportions. Youden indices for models W1, W2, and LL1 under scheme 2 are listed in Table 9 and support the results in Table 6.

**Table 9:**
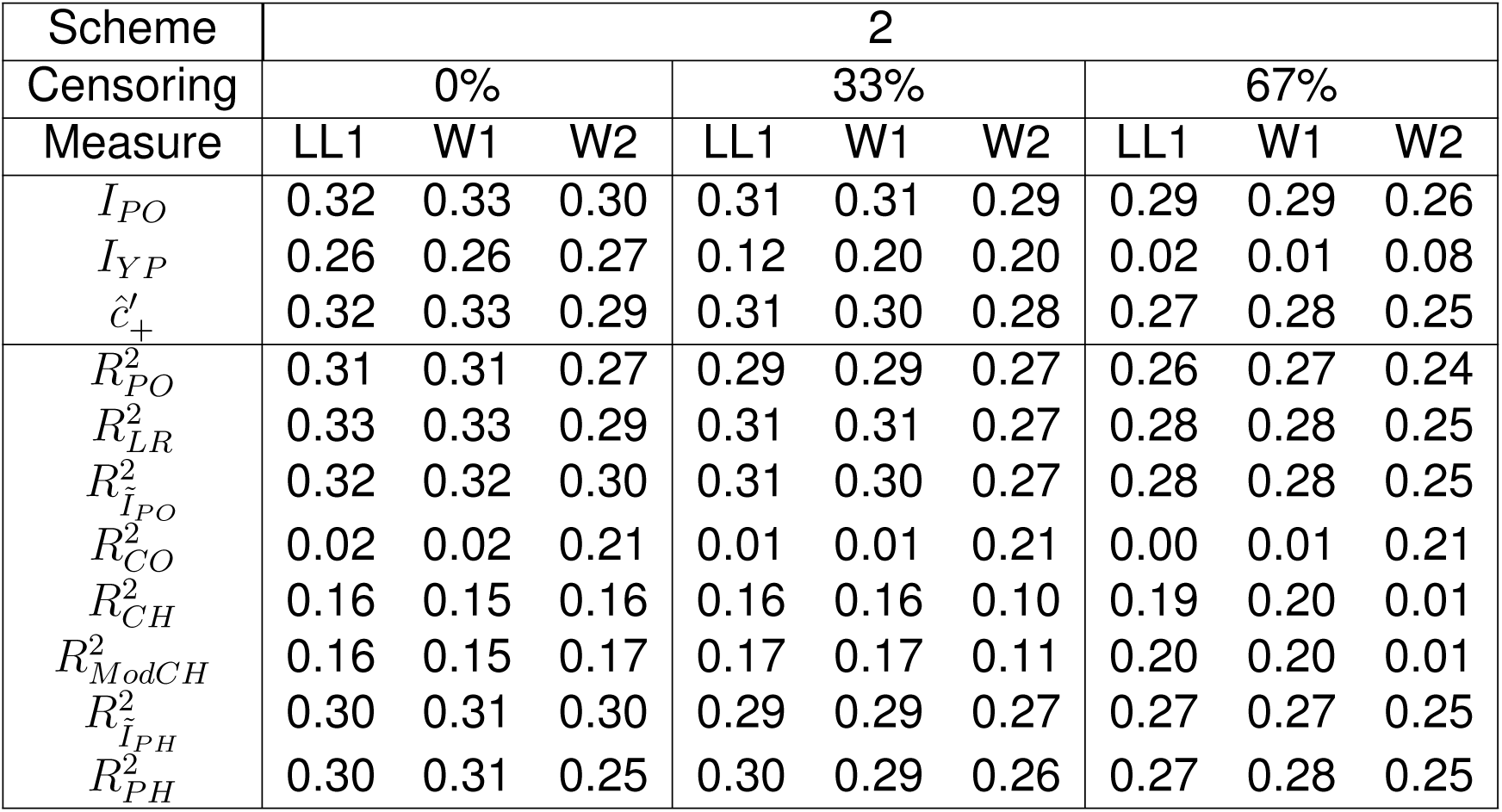
Simulation Scheme 2: Comparison of methods (Youden Index)

### 11 Application to Genomic Data

Once a subset of genomic features has been selected at a particular threshold, the combined effect of these features on survival can be evaluated using a weighted average of feature expression. Let *m* be the number of features in a given subset of interest. If ***β*** = {*β*_1_, …, *β*_*m*_}′ and **Z** are the corresponding regression coefficient vector and the *n* × *m* expression matrix for features in this subset, then a weighted average, ***η***, can be calculated as ***η*** = **Z*β***. Thus, ***η*** is a vector of size *n*, where each subject has a weighted average computed across all *m* features in a subset. This weighted average uses every feature in each subset where each features’s contribution is quantified by the estimate of the coefficient. It is worth noting that the linear predictor ***η*** can be interpreted as the logarithm of the hazard ratio for the PH model and as the logarithm of the odds ratio for the PO model. A graphical analysis of the combined effect of features selected using 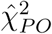 (as shown in Table 4) was performed for representative data sets from different data types. This included data sets 1 (methylation), 2 (microRNA expression), 3 (RNA sequencing), 4 (mRNA expression) and 8 (CNV). Panels (a), (b) and (c) in Figures 8-12 represent KM survival curves, cumulative hazard curves (on the log-scale) and odds curves, respectively, for subjects with high and low weighted average feature expression (determined by the median split) for each of these data sets. We observed that when 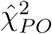 was used to select features, the PO model generally provided a good fit while the PH did not fit in some cases (as seen in panels (a), (b) and (c) of Figures 8-12). These observations were further corroborated by GOF tests for the PH, PO and YP models where the PO and YP models were found to provide a good fit for weighted feature expression in all cases. This analysis emphasizes the versatility and modeling flexibility provided by the PO and YP models in allowing PH as well as certain forms of NPH.

